# The bacterial actin-like cell division protein FtsA forms curved antiparallel double filaments upon binding of FtsN

**DOI:** 10.1101/2021.11.08.467742

**Authors:** Tim Nierhaus, Stephen H McLaughlin, Frank Bürmann, Danguole Kureisaite-Ciziene, Sarah Maslen, Mark J Skehel, Conny WH Yu, Stefan MV Freund, Louise FH Funke, Jason W Chin, Jan Löwe

## Abstract

Cell growth and division of walled bacteria depend on the synthesis and remodelling of peptidoglycan (PG). These activities are carried out by two multiprotein complexes, the elongasome and the divisome during cell elongation and division, respectively. Filaments of tubulin-like FtsZ form the cytoplasmic scaffold for divisome assembly, the Z-ring. In *E. coli*, the actin homologue FtsA anchors the Z-ring to the membrane and recruits downstream divisome components, including bitopic FtsN. FtsN is recruited late and activates the periplasmic PG synthase FtsWI. To start unravelling the activation mechanism involving FtsA and FtsN, we showed that *E. coli* FtsA forms antiparallel double filaments on lipid monolayers when also binding FtsN’s cytoplasmic tail, and that *Vibrio maritimus* FtsA crystallised as an equivalent double filament. We structurally located the FtsA-FtsN interaction site in FtsA’s IA-IC interdomain cleft and confirmed FtsA double filament formation *in vivo* using site-specific cysteine cross-linking. FtsA-FtsN double filaments reconstituted on and in liposomes preferred negative Gaussian curvature, as was previously shown for the elongasome’s actin, MreB. MreB filaments serve as curvature-sensing “rudders”, orienting insertion of PG around the cell’s circumference. We propose that curved antiparallel FtsA double filaments function similarly in the divisome: FtsA filaments, together with dynamic FtsZ filaments orient and concentrate cell-constricting septal PG synthesis in the division plane.

## INTRODUCTION

In non-spherical, walled bacteria, cell shape is determined by the peptidoglycan sacculus, the load-bearing structure counteracting turgor pressure (Typas *et al.*, 2011). Insertion of new glycan strands into the sacculus is orchestrated by two related multiprotein complexes: the divisome, acting in cell division, and the elongasome (or rod complex), acting in cell elongation (Szwedziak and Löwe, 2013). In *Escherichia coli*, both complexes span the entire cell envelope and contain the cytoplasmic, membrane-binding and filament-forming actin homologues FtsA (divisome) and MreB (elongasome) (Pichoff and Lutkenhaus, 2005; Salje *et al.*, 2011).

In *E. coli* and many other bacteria, FtsA is the main membrane anchor for the cell division ring, the Z-ring, which initiates and organises cell division (Geissler *et al.*, 2003; Pichoff and Lutkenhaus, 2002). The Z-ring is mainly formed by filaments of the tubulin homologue FtsZ (Bisson-Filho *et al.*, 2017; Nogales *et al.*, 1998; Yang *et al.*, 2017). FtsZ treadmilling dynamics, driven by FtsZ’s GTPase activity, were shown to be essential for the initial condensation of FtsZ filaments into the Z-ring, but seem to become dispensable after partial constriction of the septum (Monteiro *et al.*, 2018; Squyres *et al.*, 2021; Whitley *et al.*, 2021).

Apart from localising FtsZ to the membrane, FtsA is also involved in the recruitment of downstream divisome components such as FtsK, FtsN and potentially FtsQ (Du and Lutkenhaus, 2017; Karimova *et al.*, 2005). Together with the bitopic membrane protein FtsN, FtsA is required for divisome integrity (Pichoff *et al.*, 2018). FtsN assembles last into the divisome and activates the bipartite septal PG synthase FtsWI via FtsQLB (Gerding *et al.*, 2009; Li *et al.*, 2021; Liu *et al.*, 2015; Marmont and Bernhardt, 2020; Tsang and Bernhardt, 2015), and possibly also PBP1b (Boes *et al.*, 2019; Müller *et al.*, 2007).

Small amounts of FtsN are recruited early to the divisome in a FtsA-dependent manner (Busiek and Margolin, 2014). The FtsA-FtsN interaction is known to involve FtsA’s for actin-like proteins uniquely positioned IC domain and FtsN’s cytoplasmic tail, which comprises ∼32 amino acids in *E. coli* (Baranova *et al.*, 2020; Busiek *et al.*, 2012; Pichoff *et al.*, 2015). Two FtsN suppressor mutations in FtsA located in its IC domain (Bernard *et al.*, 2007; Liu *et al.*, 2015) further support the idea that FtsN binds the IC domain of FtsA. In FtsN, a conserved stretch of basic amino acids in its cytoplasmic tail is required for interaction with FtsA *in vitro* (Baranova *et al.*, 2020; Busiek *et al.*, 2012). In a *zipA* null background, in which the FtsA-FtsN interaction becomes essential, mutation of the basic stretch in FtsN was however permissible, whereas a D5N mutation was not (Pichoff *et al.*, 2015). It has been proposed that interaction with FtsN depolymerises FtsA and, thereby, allows recruitment of downstream divisome components via binding sites that were (partially) occluded in the FtsA polymer (Pichoff *et al.*, 2015).

However, little else has been reported regarding FtsA’s polymerisation state in the divisome and its interaction sites with other divisome proteins. Despite the uniqueness of FtsA’s IC domain amongst the actin-like proteins (van den Ent and Löwe, 2000), FtsA was shown to form single protofilaments that recapitulate some structural features of *bona fide* actin protofilaments (Szwedziak *et al.*, 2012). Double filaments were shown to be the smallest functional unit of all other known actin homologues (Wagstaff and Löwe, 2018) and, indeed, *Thermotoga maritima* FtsA was shown to form double filaments (Szwedziak *et al.*, 2012). Furthermore, several mutations in *E. coli* FtsA, originally described as ZipA suppressor mutations, were shown to facilitate double filament formation of FtsA on supported lipid monolayers (Schoenemann *et al.*, 2018).

In contrast, the actin-like protein of the elongasome, MreB, adopts the canonical actin-like fold (van den Ent *et al.*, 2001). MreB binds to membranes directly and forms curved antiparallel double filaments (Hussain *et al.*, 2018; Salje *et al.*, 2011; van den Ent *et al.*, 2014). This enables a curvature-sensing mechanism that allows MreB filaments to align with the axis of highest principal curvature, the short axis of the cell in the case of rod-shaped cells, such as *Bacillus subtilis* (Hussain *et al.*, 2018; Wong *et al.*, 2019). Hence, the elongasome uses MreB double filaments as “rudders” to direct movement of its bipartite PG synthase RodA-PBP2 (equivalent to FtsWI in the divisome) and, thereby, directs insertion of new PG strands around the cell circumference (Dion *et al.*, 2019; Garner *et al.*, 2011). It is thought that the radially inserted PG hoops enforce rod shape by mechanically limiting cell width expansion.

Here, we showed that FtsA from the Gram-negative bacterium *Vibrio maritimus* crystallised as antiparallel double filaments. Using supported lipid monolayers, we demonstrated that *E. coli* FtsA forms very similar antiparallel double filaments upon binding the short, cytoplasmic tail of FtsN. Using crystallography, we showed that FtsN binds FtsA in the IA-IC interdomain cleft. Furthermore, we demonstrated the *in vivo* significance of FtsA double filaments by site-specific cysteine cross-linking in *E. coli*. FtsA-FtsN filaments reconstituted on and in liposomes preferred negative Gaussian curvature, suggesting that FtsA double filaments in the divisome sense curvature using a similar mechanism as has been proposed for MreB filaments in the elongasome. We therefore propose that FtsA double filaments provide a curvature-guided mechanism to orient cell-constricting septal PG synthesis, which is further organised by dynamic FtsZ filaments into the required ring structure at the division site.

## RESULTS

### *Vibrio maritimus* FtsA crystallises as an antiparallel double filament

Previously obtained structures of FtsA are limited to the two organisms *Staphylococcus aureus* and *Thermotoga maritima* (Fujita *et al.*, 2014; Szwedziak *et al.*, 2012; van den Ent and Löwe, 2000). Encouraged by a recent study reporting double filaments formed by *E. coli* FtsA (Schoenemann *et al.*, 2018), we reasoned that further FtsA crystal structures, especially from Gram-negative organisms such as *E. coli*, might provide new insights into the architecture of FtsA filaments and their function in the divisome.

We solved the crystal structure of C-terminally truncated FtsAs from *E. coli* (EcFtsA, Protein Data Bank (PDB) identifier: 7Q6D) and the closely related Gram-negative bacterium *Xenorhabdus poinarii* (PDB 7Q6G, 90 % identity of XpFtsA to EcFtsA) and *Vibrio maritimus* (PDB 7Q6F, 70 % identity of VmFtsA to EcFtsA) (Supplementary Table T 1). The EcFtsA^1-405^ and XpFtsA^1-396^ structures showed single protofilaments with a loose longitudinal contact between the IIA and IIB subdomains (Supplementary Figure S1a). In contrast, VmFtsA^1-396^ crystallised as straight, antiparallel double filaments, with lateral contacts formed by the IC domain, which among actins is unique to FtsA (van den Ent and Löwe, 2000) (Figure 1a). The longitudinal interface contact between the IIA-IIB domains was tight in both protofilaments, as was the case for TmFtsA bound to ATP-γ-S (PDB 4A2B) (Szwedziak *et al.*, 2012) (Supplementary Figure S1a). We noticed that the position of the IC domain is variable within the FtsA monomers across FtsA structures (Supplementary Figure S1b). Positioning of the IC domain did neither correlate with species nor the presence of continuous protofilaments within the crystal, as analysed for PDB 1E4F, 4A2B and 7Q6F.

**Figure 1.**
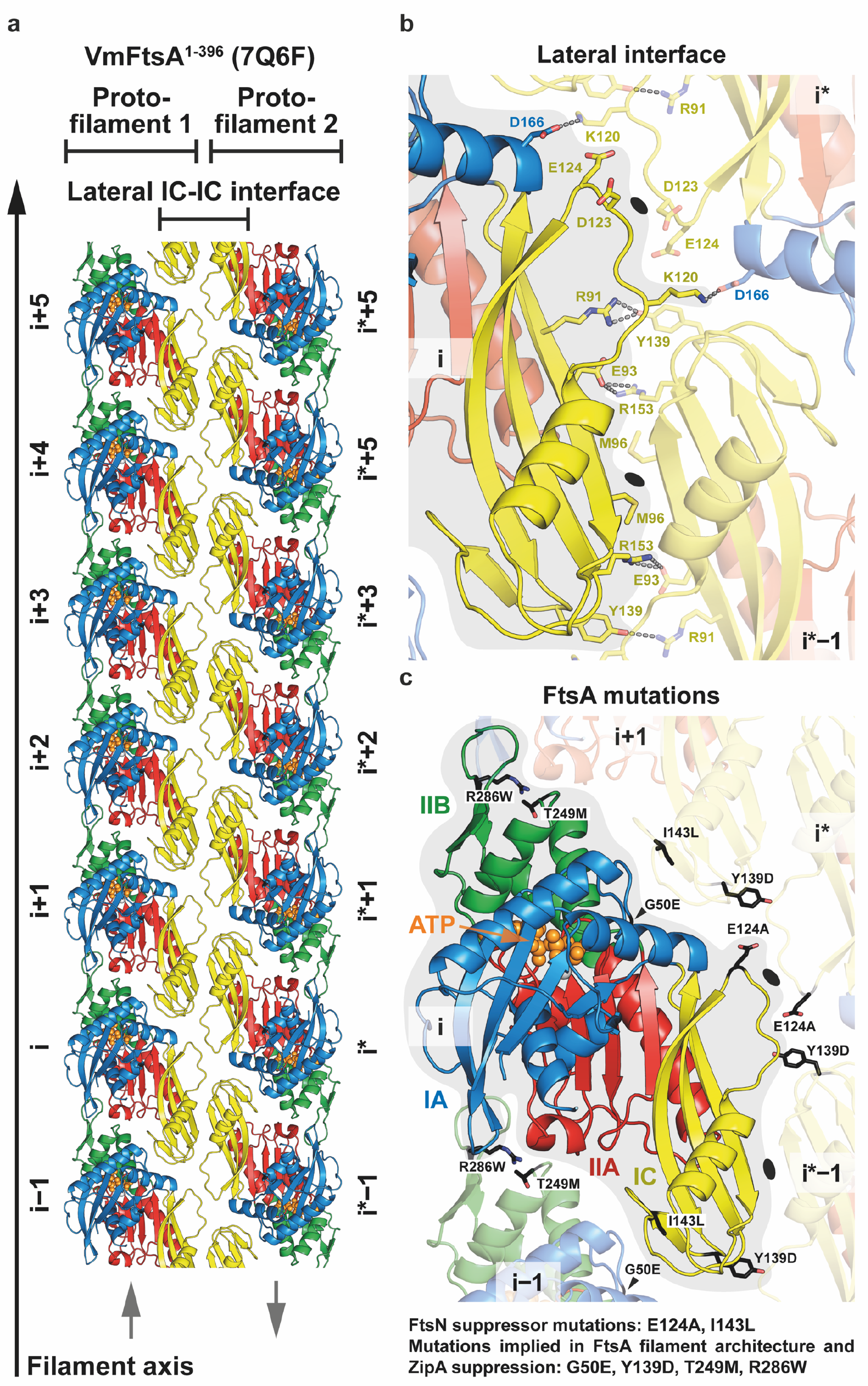
*Vibrio maritimus* FtsA crystallises as antiparallel double filaments via lateral contacts between IC domains. **a**, Top view of the VmFtsA^1-396^ antiparallel double filament in cartoon representation from the membrane-proximal side (PDB 7Q6F). The lateral interface is almost exclusively formed by the IC domain (yellow). Grey arrows indicate the relative orientations of FtsA monomers in each protofilament. **b**, Side chain interactions in the lateral filament interface. Each FtsA_i_ contacts two neighbouring FtsAs, FtsA_i-1*_ and FtsA_i*_, in the opposing protofilament. Both interfaces have local C2 symmetry (black ellipses). **c**, Reported FtsN suppressor mutations in *E. coli* FtsA, E124A (Bernard *et al.*, 2007) and I143L (Liu *et al.*, 2015), and ZipA suppressor mutations that recently have been implied in FtsA (double) filament formation (Schoenemann *et al.*, 2018) are mapped on the VmFtsA double filament structure. All mutations are part of filament interfaces.

The lateral interface in the VmFtsA double filament is almost exclusively formed by contacts between IC domains (Figure 1b). Each FtsA monomer contacts two subunits in the opposing protofilament, forming the FtsA_i_-FtsA_i*_ and FtsA_i_-FtsA_i*−1_ lateral interfaces. Both interfaces have local twofold (C2) symmetry as indicated in Figure 1b. The VmFtsA double filament is wider than the MreB double filament (PDB 4CZJ) (van den Ent *et al.*, 2014), which is, like FtsA filaments, comprised of two antiparallel protofilaments (Supplementary Figure S1c). The membrane-proximal side of FtsA and MreB double filaments is flat. ZipA suppressor mutations in *E. coli* FtsA (including FtsA^R286W^, also known as FtsA*), which were described to also affect filament architecture (Schoenemann *et al.*, 2018), correspond to residues in subunit interfaces in the VmFtsA double filament structure (Figure 1c). Interestingly, the FtsN suppressor mutations E124A (Bernard *et al.*, 2007) and I143L (Liu *et al.*, 2015) described for *E. coli* FtsA also map to subunit interfaces in the VmFtsA double filament structure (Figure 1c).

### *E. coli* FtsA forms antiparallel double filaments upon binding the cytoplasmic tail of FtsN, EcFtsN^1-32^

As described previously (Krupka *et al.*, 2017), we found that *E. coli* FtsA forms “mini-rings” on supported lipid monolayers (Figure 2g, “WT + no peptide”). Given that filament architecture mutations and FtsN suppressor mutations in *E. coli* FtsA map to lateral interfaces in the VmFtsA double filament (Figure 1c), we hypothesised that FtsN could induce double filament formation of *E. coli* FtsA. The short, cytoplasmic tail of FtsN (comprising ∼32 amino acids in *E. coli*) (Supplementary Figure S2f) interacted with FtsA as had been shown previously (Busiek *et al.*, 2012; Busiek and Margolin, 2014; Pichoff *et al.*, 2015). Using surface plasmon resonance (SPR), we showed that EcFtsN^1-32^ binds EcFtsA and EcFtsA^1-405^, a C-terminal truncation of EcFtsA lacking the amphipathic membrane binding helix, with dissociation constants (K_d_) of 0.8 µM and 2.0 µM, respectively (Figure 2a). The interaction between *V. maritimus* FtsA^1-396^ and FtsN^1-29^ was about three-fold weaker than the EcFtsA^1-405^-EcFtsN^1-32^ interaction (Supplementary Figure S2a). Since cross-linking of FtsA to the flow cell surface in SPR might affect FtsA polymerisation, we also probed the FtsA-FtsN interaction in solution using fluorescence polarisation (FP), for which C-terminally truncated FtsA was titrated into fluorescently labelled FtsN peptide. FP data were fitted to a two-step model with K_d_s of about 0.016 µM and 11 µM for the EcFtsA^1-405^-FtsN^1-32-C-Atto 495^ interaction (Figure 2b). Again, the VmFtsA^1-396^-FtsN^1-29^ interaction was about four-fold weaker for the first binding event (Supplementary Figure S2b). Next, we subjected the system to analytical ultracentrifugation using a fluorescence detection system (FDS-AUC) to characterise the two binding events of the FtsA-FtsN interaction. FDS-AUC showed that both binding events are accompanied by formation of higher order FtsA-FtsN aggregates or polymers, indicating that the cytoplasmic FtsN peptide not only binds to FtsA but could also facilitate FtsA polymerisation (Figure 2c; Supplementary Figure S2c). Using a co-pelleting assay, for which cytoplasmic FtsN peptide was titrated into truncated FtsA, we showed that FtsN peptide indeed induces FtsA polymerisation, as apparent by the increased fraction of FtsA in the pellet at higher concentrations of FtsN peptide (Figure 2d; Supplementary Figure S2d).

**Figure 2:**
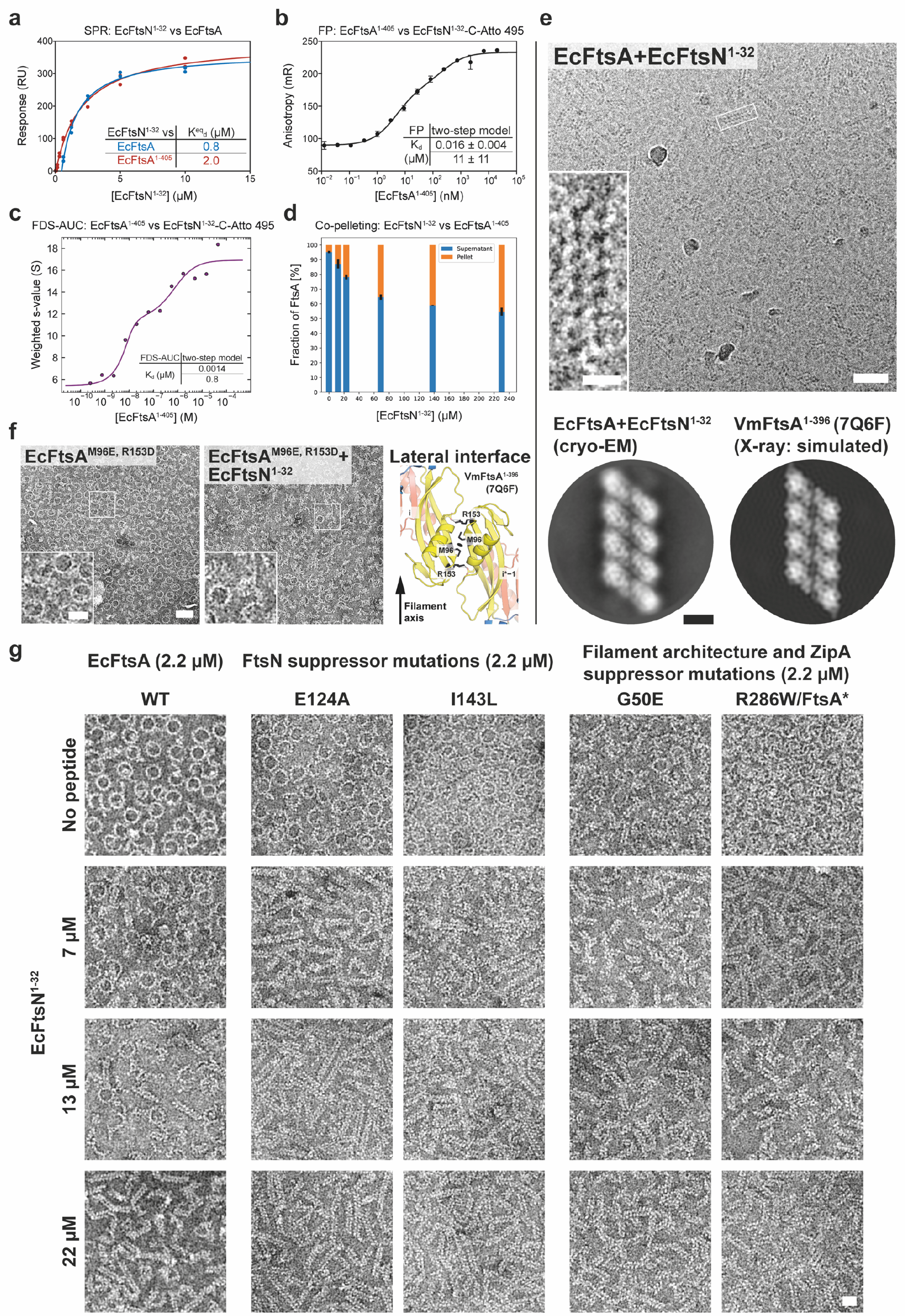
*E. coli* FtsA forms antiparallel double filaments upon binding the cytoplasmic tail of FtsN, EcFtsN^1-32^. **a**, SPR equilibrium response titration of EcFtsN^1-32^ binding to immobilised EcFtsA (blue) or EcFtsA lacking its C-terminal membrane binding helix, EcFtsA^1-405^ (red). EcFtsN^1-32^ has micromolar affinity for both FtsA proteins. **b**, EcFtsA^1-405^ titration into fluorescently labelled EcFtsN^1-32^-Cys-Atto 495. Data were fitted with a two-step model, with transitions being indicative of FtsN binding and polymerisation (panel c). K_d_s are given as mean ± SEM. **C**, Change in the weight-averaged sedimentation coefficient of a EcFtsA^1-405^ titration into fluorescently labelled EcFtsN^1-32^-Cys-Atto 495 by fluorescence detection system (FDS)-AUC shows that EcFtsN^1-32^ is part of higher order FtsA polymers. Data were fitted to a two-step model, recapitulating the FP data in panel b. **d**, Co-pelleting assay of EcFtsN^1-32^ titrated into EcFtsA^1-405^. The fraction of FtsA in the pellet (orange) increases with increasing amounts of EcFtsN^1-32^, indicating that EcFtsN^1-32^ induces FtsA polymerisation. Given are mean ± sd (black lines) of technical duplicates. **e**, EcFtsA and EcFtsN^1-32^ form double filaments on supported lipid monolayers as determined by cryo-EM. 2D classification confirmed the antiparallel arrangement of the observed double filaments. A computed 2D projection from the VmFtsA double filament crystal structure (PDB 7Q6F) is shown for comparison. Scale bars, 50 nm, 10 nm (inset), 5 nm (2D class averages). **f**, Based on the VmFtsA crystal structure (PDB 7Q6F), a lateral interface mutant of *E. coli* FtsA, EcFtsA^M96E, R153D^, was designed. EcFtsA^M96E, R153D^ is deficient in FtsN^1-32^-dependent double filament formation as determined by negative stain electron microscopy on a supported lipid monolayer. Scale bar, 50 nm, 20 nm (inset). **g**, Negative stain electron micrographs of EcFtsA and EcFtsA mutants on supported lipid monolayers with increasing concentrations of EcFtsN^1-32^. EcFtsA forms “mini-rings” in the absence of FtsN peptide, as described previously (Krupka *et al.*, 2017). With increasing EcFtsN^1-32^ concentrations, EcFtsA forms fewer “mini-rings” and more double filaments, until only double filaments are present at ten-fold molar excess of EcFtsN^1-32^. The FtsN suppressor mutants FtsA^E124A^ and FtsA^I143L^ more readily form double filaments at lower EcFtsN^1-32^ concentrations than wildtype FtsA. Introduction of the G50E or R286W mutations into EcFtsA decreases the number of “mini-rings” formed in the absence of EcFtsN^1-32^. Both mutations also facilitate FtsA double filament formation at lower concentrations of EcFtsN^1-32^, compared to wildtype. See Figure 1c for the position of the mutations in the double FtsA filament. Scale bar, 20 nm.

Given that the cytoplasmic peptide of FtsN induces FtsA polymerisation, we next investigated the architecture of those polymers using negative stain electron microscopy (EM). Both EcFtsA-FtsN^1-32^ and VmFtsA-FtsN^1-29^ formed short double filaments on supported lipid monolayers (Figure 2g, “WT”; Supplementary Figure S2e). Electron cryo-microscopy (cryo-EM) imaging and subsequent 2D averaging of EcFtsA-FtsN^1-32^ double filaments on lipid monolayers showed that EcFtsA-FtsN^1-32^ double filaments have very similar architecture to the VmFtsA^1-396^ double filaments determined by crystallography (PDB 7Q6F) (Figure 2e). To corroborate, we designed a lateral interface mutant of *E. coli* FtsA based on the VmFtsA double filament structure: EcFtsA^M96E, R153D^ (Figure 2f, right). The lipid monolayer assay showed that EcFtsA^M96E, R153D^ was indeed deficient in EcFtsN^1-32^-dependent double filament formation (Figure 2f).

The short cytoplasmic tail of FtsN harbours two sequence motifs that are conserved among *E. coli* and related proteobacteria with similar FtsA and FtsN sequences: a conserved ^3^R/KDY^6^ (*E. coli* amino acid positions) motif near the N-terminus (Pichoff *et al.*, 2015) and two to three stretches of basic amino acids, the most prominent of which in *E. coli* FtsN is ^16^RRKK^19^ (Busiek *et al.*, 2012) (Supplementary Figure S2f, g). In *E. coli*, a D5N mutation in the conserved N-terminal motif of FtsN was shown to impair FtsA-FtsN interactions *in vivo* (Pichoff *et al.*, 2015), but did not affect binding or co-localisation with FtsA-FtsZ filaments on supported lipid bilayers *in vitro* (Baranova *et al.*, 2020). In contrast, and somewhat confusingly, mutation of the basic ^16^RRKK^19^ stretch abrogated binding *in vitro* (Baranova *et al.*, 2020; Busiek *et al.*, 2012), yet was permissible *in vivo* under conditions for which the FtsA-FtsN interaction becomes essential (Pichoff *et al.*, 2015). We therefore tested a set of EcFtsN^1-32^ truncations and mutations in our SPR and monolayer assays to determine the effect of those modifications on FtsA-FtsN interactions and on double filament formation (Supplementary Figure S2h, i). Mutation or deletion of any one of the three stretches of basic amino acids in EcFtsN^1-32^ reduced binding affinity for FtsA and, consequently, its ability to promote FtsA double filament formation. EcFtsN^1-32, D5N^ and EcFtsN^1-33, scrambled^ (Supplementary Table T 4), which has been described previously but was shortened through removal of its tags in this study (Baranova *et al.*, 2020), did not show reduced binding affinity. But, EcFtsN^1-32, D5N^ and EcFtsN^1-33, scrambled^ were unable to induce FtsA double filament formation. Hence, the D5N mutation in FtsN might be non-permissible *in vivo* under conditions for which the FtsA-FtsN interaction becomes essential because it fails to promote formation of FtsA double filaments. In other words, this result supports the idea that FtsA double filaments are important for the activation of cell division by FtsA and FtsN.

We then tested whether any of the FtsN suppressor (Bernard *et al.*, 2007; Liu *et al.*, 2015) or filament architecture and ZipA suppressor mutations (Schoenemann *et al.*, 2018) in FtsA would impact FtsN-induced double filament formation of FtsA (Figure 1c). While the FtsN suppressor mutants FtsA^E124A^ and FtsA^I143L^, similar to wildtype FtsA, formed “mini-rings” in the absence of EcFtsN^1-32^, they more readily formed double filaments than wildtype FtsA at three-(7 µM) and six-fold (13 µM) molar excess of EcFtsN^1-32^ (Figure 2g). The filament architecture and ZipA suppressor mutants FtsA^G50E^ and FtsA^R286W^ (also known as FtsA*) formed fewer “mini-rings” than wildtype FtsA in the absence of FtsN (Figure 2g). For FtsA^G50E^, “mini-rings” and short double filaments were observed. FtsA^R286W^ predominantly formed arcs, curved, short and single filaments. Similar to the FtsA^E124A^ and FtsA^I143L^ proteins, FtsA^G50E^ and FtsA^R286W^ formed many more double filaments than wildtype FtsA at three-(7 µM) and six-fold (13 µM) molar excess of EcFtsN^1-32^ (Figure 2g). Thus, both FtsN suppressor as well as ZipA suppressor and FtsA filament architecture mutants appear to facilitate FtsN-dependent FtsA double filament formation.

### VmFtsN^1-29^ binds VmFtsA^1-396^ in the IA-IC interdomain cleft

To understand the FtsA-FtsN interaction at the molecular level, we co-crystallised VmFtsA^1-396^ with the *V. maritimus* cytoplasmic FtsN peptide, VmFtsN^1-29^. Although similar crystallisation conditions were used, the VmFtsA^1-396^-VmFtsN^1-29^ co-crystal structure (PDB 7Q6I) showed a larger unit cell than that obtained for the VmFtsA^1-396^ double filament structure [PDB 7Q6F, two FtsA monomers per asymmetric unit (ASU)], encompassing 16 FtsA monomers per ASU (Supplementary Table T 1). The 16 FtsA monomers are organised into four tetramers, each containing two antiparallel very short protofilaments, each comprised of two FtsA monomers (Figure 3a). One of the two FtsA monomers in a protofilament adopts a closed and the other one an open conformation (Supplementary Figure S3a). The positions of the IC domain in the closed (PDB 7Q6I chain A) and open (PDB 7Q6I chain B) conformation are similar but not identical to the IC domain position in the VmFtsA^1-396^ double filament (PDB 7Q6F) (Supplementary Figure S1b). Only in the IA-IC interdomain cleft of closed FtsA monomers could we find electron density for the VmFtsN^1-29^ peptide (Figure 3a, c; Supplementary Figure S3a). The IC domain in the open conformation is rotated downwards by 13.8 ° compared to the closed conformation, as analysed by DynDom (Hayward and Lee, 2002) (Supplementary Figure S3a). This means the open conformation is likely incompatible with VmFtsN^1-29^ binding (Supplementary Figure S3a). The IC domains in the lateral interface of each tetramer are twisted against each other compared to the straight VmFtsA^1-396^ double filament (Figure 3a). Several hydrogen bonding contacts are lost because of the bending when compared to the straight filaments (Figure 3a, asterisks; Figure 3b). The VmFtsA^1-396^-VmFtsN^1-29^ tetramer is bent along the filament axis with an estimated radius of curvature of 16.3 nm (Figure 3b).

**Figure 3.**
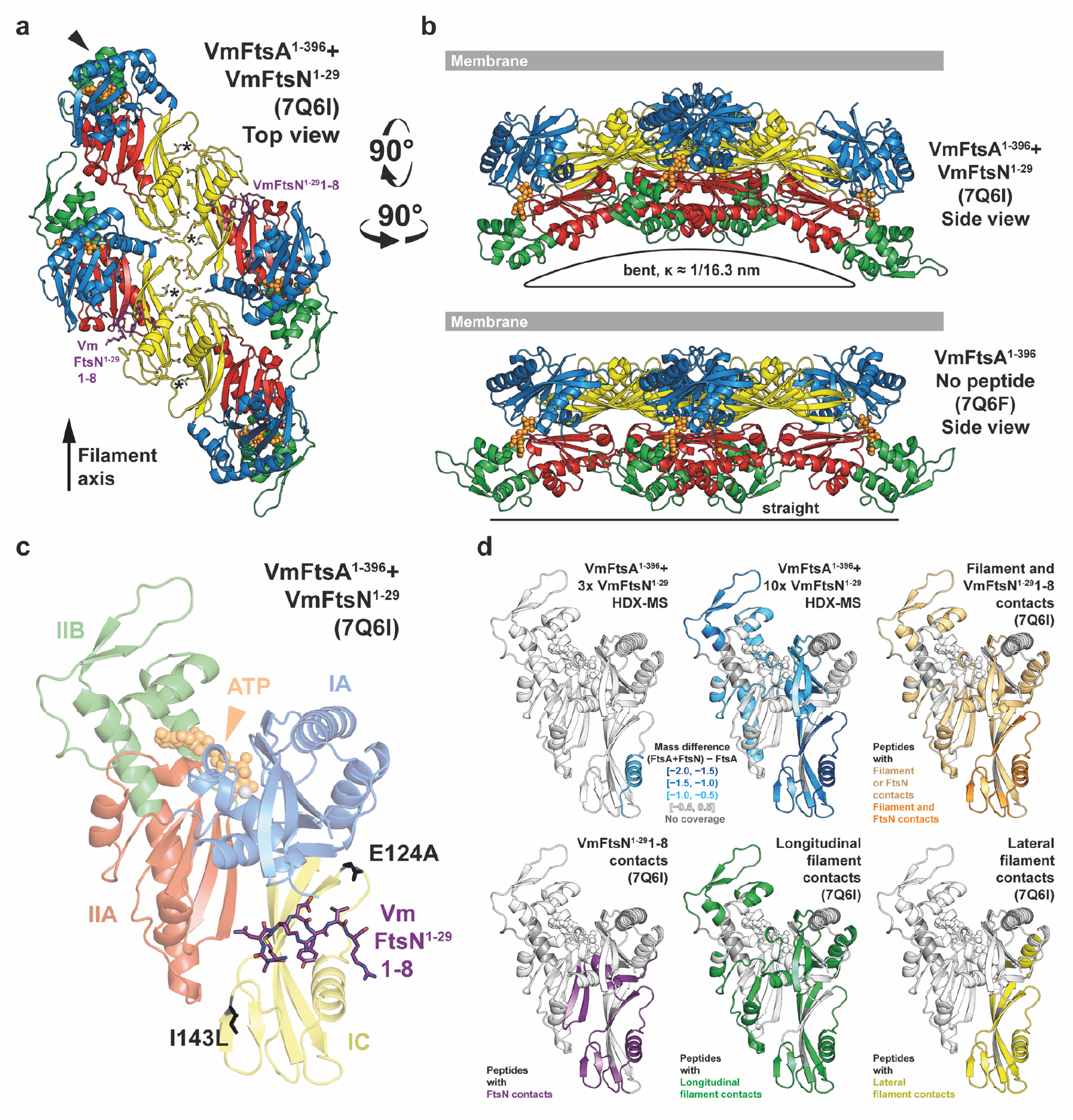
VmFtsA^1-396^ and VmFtsN^1-29^ co-crystallise as short, bent double filaments, with VmFtsN^1-29^ binding FtsA in the IA-IC interdomain cleft. **a,** 16 FtsA monomers in the ASU of the co-crystal structure are organised into short, antiparallel, and curved tetramers. Top view of a representative tetramer (which requires crystal symmetry to be applied to the PDB coordinates) from the membrane-proximal side. Residues 1-8 of VmFtsN^1-29^ (purple sticks) are present in the closed subunit of each protofilament. IC domains are tilted against each other compared to the VmFtsA^1-396^ double filament structure (PDB 7Q6F), presumably destabilising the R91-Y139 interaction (asterisks) in the lateral interface. The IIB domain of the top left monomer (chain D) is partially disordered (arrowhead). **b**, A comparison of the side views of the PDB 7Q6I and PDB 7Q6F crystal structures illustrates that the 7Q6I tetramer is bent along the filament axis. **c**, Residues 1-8 of VmFtsN^1-29^ bind in the IA-IC interdomain cleft of VmFtsA^1-396^. Positions of FtsN suppressor mutations E124A (Bernard *et al.*, 2007) and I143L (Liu *et al.*, 2015) are shown in black. **d**, HDX-MS analysis of VmFtsA^1-396^ with three-(30 µM) or ten-fold molar excess (100 µM) of VmFtsN^1-29^, confirming VmFtsN^1-29^ to bind in the IA-IC interdomain cleft of FtsA. Peptides that are protected in the presence of VmFtsN^1-29^ (slower exchange of hydrogen for deuterium) are highlighted in blue. Presumably because high concentrations of VmFtsN^1-29^ induce polymerisation of VmFtsA^1-396^ (Supplementary Figure S2c, d), peptides mapping to the filament interfaces are protected as well. For orientation, peptides are coloured according to their involvement in different interfaces of the VmFtsA^1-396^-VmFtsN^1-29^ co-polymer (PDB 7Q6I).

Due to the low resolution (3.6 Å) of the VmFtsA^1-396^-VmFtsN^1-29^ co-crystal structure (Supplementary Table T 1), we could not unambiguously interpret the electron density map representing about seven to eight amino acids of the VmFtsN^1-29^ peptide. We therefore confirmed the location of the VmFtsN^1-29^ binding site in VmFtsA^1-396^ by hydrogen deuterium exchange mass spectrometry (HDX-MS) (Figure 3d). At three-fold molar excess (30 µM) of VmFtsN^1-29^ over VmFtsA^1-396^, only helix 2 located in the IC domain of FtsA showed significantly reduced deuterium incorporation (lower mass differential) compared to free VmFtsA^1-396^ (Figure 3d, top left). This agrees with VmFtsN^1-29^ binding across the IC domain as seen in the co-crystal structure (Figure 3c). At 10-fold molar excess (100 µM) of VmFtsN^1-29^, FtsA peptides in filament interfaces were also protected, again, indicating that binding of the cytoplasmic FtsN peptide to FtsA facilitates filament formation (Figure 3d).

The FtsA-FtsN interaction is reminiscent of that between PilM and PilN, two proteins involved in type IV pilus formation (Karuppiah and Derrick, 2011; McCallum *et al.*, 2016). PilM, which also has a IC subdomain, is a structural homologue of FtsA, and PilN, similar to FtsN, is a bitopic membrane protein with a short, cytoplasmic tail comprising about 30 amino acids (Karuppiah and Derrick, 2011; McCallum *et al.*, 2016). FtsN and PilN both bind in the IA-IC interdomain cleft (assuming FtsA domain nomenclature for PilM), but use somewhat different binding modes (Supplementary Figure S3b). PilN makes extensive contacts with the IA domain of PilM. FtsN binds further downward in the cleft, predominantly contacting the IC domain of FtsA.

As mentioned, the low resolution of the electron density for the VmFtsN^1-29^ peptide in the VmFtsA^1-396^-VmFtsN^1-29^ co-crystal structure limited our ability to assign the peptide register and only peptide density associated with two FtsA monomers (PDB 7Q6I chains A and C) was interpretable. Based on the geometry *in vivo*, where FtsA binds to the cell membrane through its C-terminal amphipathic helix, we reasoned that the N-terminal half of VmFtsN^1-29^ is expected to interact with FtsA, as the globular domain of FtsA was reported to be several nanometres away from the inner membrane in *E. coli* (Szwedziak *et al.*, 2014). The C-terminal half of the VmFtsN^1-29^ peptide would be confined to the proximity of the inner membrane, as it is linked to the transmembrane helix in the full-length FtsN protein (Supplementary Figure S2f). Amino acids M1-R8 of VmFtsN^1-29^ provided the best fit for the density with reasonable chemistry, i.e. several hydrogen bonds to FtsA and a hydrophobic pocket accommodating the central tyrosine (Y6) in the sequence stretch (Supplementary Figure S3e, left). This model also implies a central position of aspartate D5 within the FtsA-FtsN interaction site that could point to a mechanism by which the D5N mutation impairs double filament formation (Supplementary Figure S2h; Supplementary Figure S3e, left). In that case, FtsN^D5N^ would not be able to bridge FtsA’s IA and IC domain. We further studied the VmFtsA^1-396^-VmFtsN^1-29^ interaction by nuclear magnetic resonance (NMR) spectroscopy. We assigned the peptide backbone of Gly-Gly-VmFtsN^2-29^, a VmFtsN^1-29^ construct in which two glycine residues replace the N-terminal methionine (Supplementary Figure S3c). Next, we analysed peak broadening (reduction in peak intensity due to the slower tumbling rate of FtsA-bound Gly-Gly-VmFtsN^2-29^ peptide) upon addition of equimolar amounts of VmFtsA^1-396^ (Supplementary Figure S3d). Intensity reduction was most prominent in the stretch of amino acids Y6-G10 (Supplementary Figure S3d), suggesting a slightly shifted binding motif compared to our density-based assignment (M1-R8) (Supplementary Figure S3e). We were not able to generate a good fit of these residues into the electron density map of the VmFtsA^1-396^-VmFtsN^1-29^ co-crystal structure. For technical reasons, the NMR experiments were carried out at pH 6.0, whereas binding experiments were done at pH 7.7 and crystallisation was achieved at pH 8.5.

### FtsA double filaments prefer negative Gaussian curvature

Based on the observation that VmFtsA^1-396^-VmFtsN^1-29^ double filaments were curved in the co-crystal structure (Figure 3b), we hypothesised that the intrinsic curvature-preference of MreB double filaments might also be a feature of FtsA-FtsN double filaments (Hussain *et al.*, 2018; Salje *et al.*, 2011). The curvature preference of MreB filaments enables a curvature-sensing mechanism that allows them to robustly align in cells with the axis of highest principal curvature, which corresponds to the short axis of rod-shaped bacteria such as *E. coli* and *B. subtilis* (Hussain *et al.*, 2018; Wong *et al.*, 2019). The elongasome uses the aligned MreB filaments to guide glycan strand synthesis around the cell’s circumference, producing radially inserted glycan strands that are believed to mechanically reinforce rod shape (Dion *et al.*, 2019; Garner *et al.*, 2011; Hussain *et al.*, 2018).

To investigate the curvature preference of FtsA filaments, we bound EcFtsA-FtsN^1-32^ double filaments on the outside of liposomes by adding the proteins to preformed liposomes made of *E. coli* polar lipid extract. Liposomes vitrified on EM grids in the absence of proteins are typically round or oval (Figure 4a, left). In contrast, liposomes coated with EcFtsA-FtsN^1-32^ double filaments regularly showed membrane indentations with negative Gaussian curvature (Figure 4a, right). We encourage the reader to revisit previously published images of MreB double filaments bound to the outside of liposomes showing a remarkably similar binding mode (Salje *et al.*, 2011; van den Ent *et al.*, 2014). To better match the geometry inside an *E. coli* cell, we then encapsulated EcFtsA-FtsN^1-32^ double filaments in liposomes by co-dilution with CHAPS-solubilised *E. coli* total lipid extract (Szwedziak *et al.*, 2014). Protein-filled liposomes were spherocylindrical (rod-shaped) or had spherocylindrical protrusions (Figure 4b). Spherocylinders were either ∼40 nm or ∼70 nm in diameter. Thicker spherocylinders contained tightly packed EcFtsA-FtsN^1-32^ double filaments that were aligned with the short axis in the cylindrical section (Figure 4b, c). We again suggest the reader to compare with published images of MreB filaments inside liposomes showing similar filament arrangements (Hussain *et al.*, 2018). EcFtsA-FtsN^1-32^ filaments were regularly absent in the hemispheres of thick spherocylinders (Figure 4b, c). If filaments were present in the hemispheres, they showed random angular orientations (Figure 4c, bottom). Rarely, we obtained liposomes with only a few EcFtsA-FtsN^1-32^ double filaments inside (Figure 4d). Again, filaments were distributed randomly in spherical sections of the liposomes but aligned with the short axis of the liposomes in more cylindrical sections. We performed 2D averaging and image classification focussed on the membrane attachment sites of the FtsA filaments in thin and thick spherocylinders. Class averages clearly illustrated the different organisation of FtsA filaments in thin and thick spherocylinders (Figure 4e). In thin spherocylinders, FtsA was organised into bent single protofilaments, suggesting that only low concentrations of FtsN^1-32^ were encapsulated in the thin spherocylinders (Figure 4e, left). In thick spherocylinders however, FtsA formed bent double filaments, as apparent by superposition of the VmFtsA^1-396^ double filament crystal structure (PDB 7Q6F) onto the 2D class average (Figure 4e, right).

**Figure 4.**
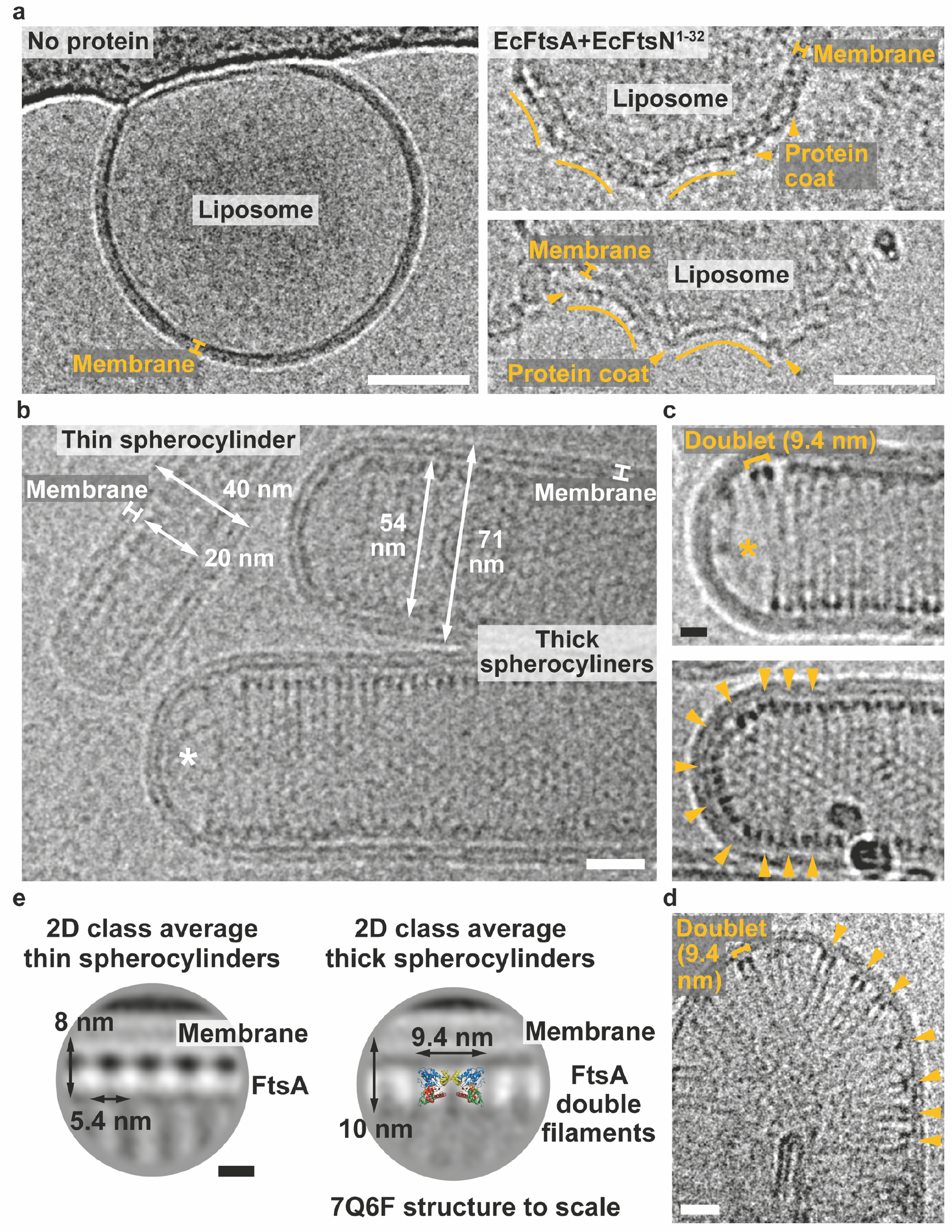
EcFtsA-FtsN^1-32^ double filaments assembled on and in liposomes show a preference for negative Gaussian curvature. **a**, Right: cryo-EM micrograph showing a spherical liposome with no proteins added. The carbon edge of the grid is visible at the top of the image. Left: cryo-EM micrographs of liposomes with EcFtsA-FtsN^1-32^ filaments (arrowheads) bound to the outside. Liposomes show indentations with negative Gaussian curvature. See Salje *et al.* for a comparison with MreB filaments added to liposomes (Salje *et al.*, 2011). Scale bars, 50 nm. **b**, Encapsulation of EcFtsA and EcFtsN^1-32^ proteins in liposomes produces thin and thick protein-filled spherocylinders (rods) with diameters of ∼40 nm and ∼70 nm, respectively. EcFtsA-FtsN^1-32^ filaments are aligned with the short axis of the spherocylinders. Note that there are no filaments in the hemisphere of a thick spherocylinder (asterisk). See Hussain *et al.* for a comparison with MreB filaments encapsulated into liposomes (Hussain *et al.*, 2018). Scale bar, 20 nm. **c**, Cryo-EM micrographs of EcFtsA-FtsN^1-32^ double filaments in thick spherocylinders. Top: Thick spherocylinders are filled with EcFtsA-FtsN^1-32^ double filaments that are aligned with the short axis. Note that there are no filaments in the hemisphere of the spherocylinder (asterisk). Bottom: EcFtsA-FtsN^1-32^ double filaments are aligned with the short axis in the cylindrical part of the spherocylinder but are randomly oriented in the hemisphere (arrowheads). Scale bar, 10 nm. **d**, Liposome with few EcFtsA-FtsN^1-32^ double filaments inside. EcFtsA-FtsN^1-32^ double filaments are randomly oriented in the semi-circular part of the liposome but are more aligned in the cylindrical part (arrowheads). Scale bar, 20 nm. **e**, 2D class averages of FtsA membrane attachment sites in thin and thick spherocylinders. EcFtsA-FtsN^1-32^ are organised into single protofilaments in thin spherocylinders, possibly due to lower amounts of EcFtsN^1-32^ being entrapped in these liposomes. Thick spherocylinders contain EcFtsA-FtsN^1-32^ double filaments. End view of the VmFtsA^1-396^ double filament structure (PDB 7Q6F) shown to scale. Scale bar, 5 nm.

### *E. coli* FtsA forms antiparallel double filaments *in vivo*

To complement and validate our *in vitro* findings, we investigated double filament formation of FtsA *in vivo* using site-specific cysteine cross-linking in *E. coli*. For greater feasibility of targeting the native *ftsA* locus, which is located in the dense *division and cell wall* (*dcw*) gene cluster, and to increase throughput with a number of mutants needed, we used a derivative of replicon <u>ex</u>cision for <u>e</u>nhanced genome engineering through programmed recombination 2 (REXER 2) (Wang *et al.*, 2016) (Supplementary Figure S4a). We inserted a *neoR* marker downstream of *lpxC* for positive selection. For visualisation via Western blotting, we introduced a 3x HA-tag (comprising 40 amino acids including linkers) into the H7-S12 loop of FtsA (Bendezu *et al.*, 2009; van den Ent and Löwe, 2000). Cysteine mutations for *in vivo* site-specific cross-linking were designed based on the VmFtsA^1-396^ double filament crystal structure (PDB 7Q6F) (Figure 5a). Mutant strains showed no growth defects and no elongated cells compared to the MG1655 parent strain (Supplementary Figure S4b-d).

**Figure 5.**
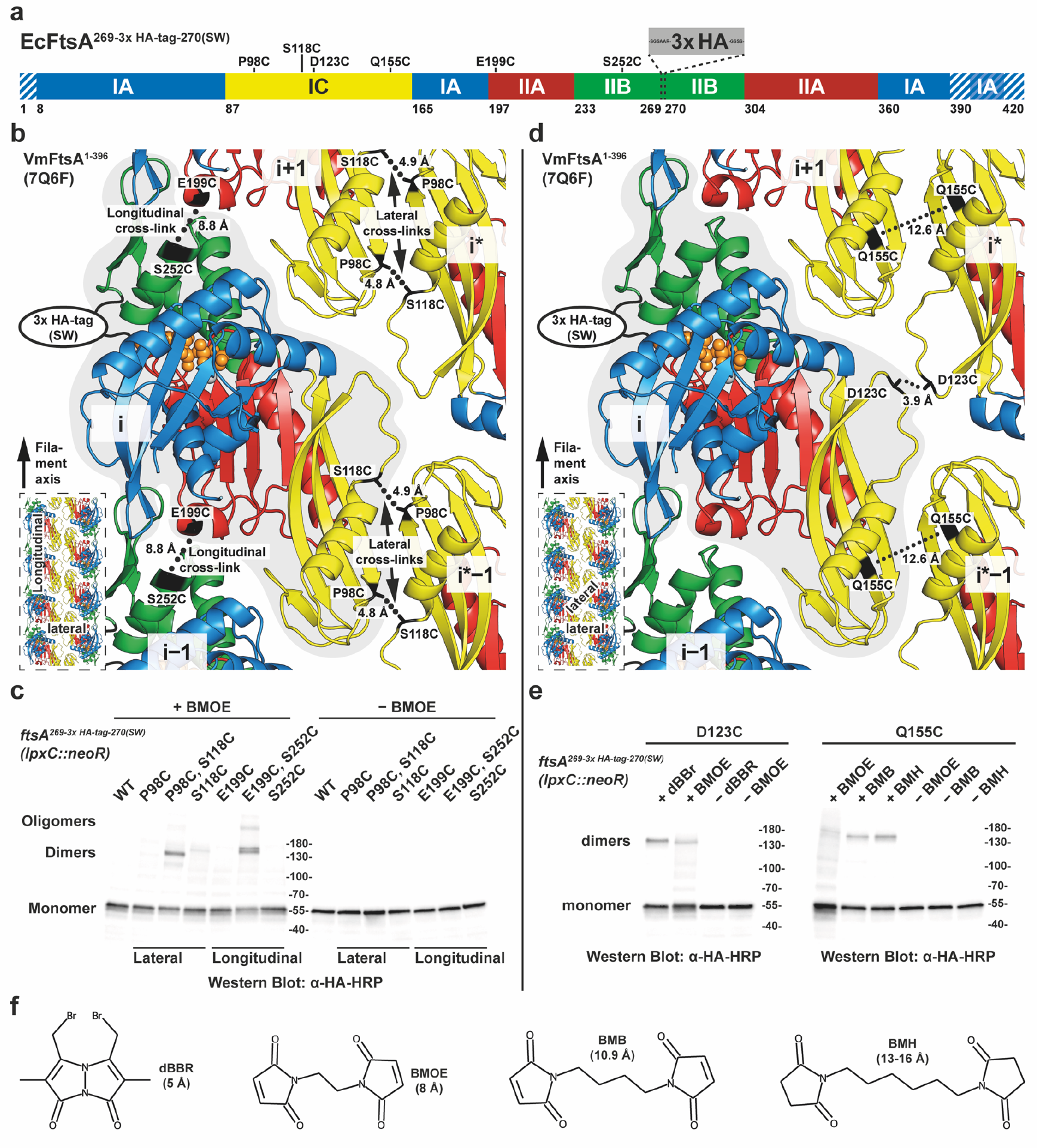
Site-specific cysteine-cross-linking in *E. coli* confirms that FtsA forms antiparallel double filaments *in vivo*. **a**, Schematic overview of the domain architecture of *E. coli* FtsA (domain boundaries and amino acids positions are the same for *V. maritimus* FtsA). Positions of cysteine mutations and the 3x HA-tag (comprising 40 amino acids including linkers, grey), which was inserted for visualisation via Western blotting, are indicated. Striped areas correspond to residues that are disordered in the VmFtsA^1-396^ double filament structure (PDB 7Q6F). **b**, Cysteine mutation pairs *ftsA^3x HA, P98C, S118C^* and *ftsA^3x HA, E199C, S252C^* probing the lateral and longitudinal filament interfaces, respectively, are highlighted on the VmFtsA double filament structure (PDB 7Q6F). C_β_-C_β_ distances between cysteines are indicated. The position of the 3x HA-tag is also shown. The inset highlights the probed interfaces in the context of the double filament. **c**, Western blot of cell lysate from FtsA cysteine mutant *E. coli* strains after *in vivo* cysteine cross-linking (+ BMOE) or without cross-linking (− BMOE). Signal for covalent FtsA dimers can be detected for the double cysteine mutants probing the lateral (*ftsA^3x HA, P98C, S118C^*) and longitudinal (*ftsA^3x HA, E199C, S252C^*) filament interfaces, but not for the respective single cysteine mutant controls. For the (*ftsA^3x HA, E199C, S252C^*) mutant, higher order oligomers can also be detected because of the open symmetry of the longitudinal contact, which leads to chaining. **d**, Cysteine mutations *ftsA^3x HA, D123C^* and *ftsA^3x HA, Q155C^* probing the lateral FtsA_i_-FtsA_i*_ and FtsA_i_-FtsA_i*−1_ filament interfaces, respectively, are highlighted on the VmFtsA double filament structure (PDB 7Q6F). C_β_-C_β_ distances between cysteines are indicated. Both single cysteine mutations utilise the local C2 symmetry of the respective lateral interface for cross-linking. The position of the 3x HA-tag is also shown. The inset highlights the probed interfaces in the context of the double filament. **e**, Western blot of cell lysate from FtsA cysteine mutant *E. coli* strains after *in vivo* cysteine cross-linking with thiol-directed cross-linkers of different lengths (“+ cross-linker”) or without cross-linking (“− cross-linker”). Signal for a FtsA dimer can be detected after using dBBr and, to a lesser extent, BMOE cross-linking in the *ftsA^3x HA, D123C^* mutant. For the *ftsA^3x HA, Q155C^* mutation (C_β_-C_β_ distance 12.6 Å), FtsA dimers can be detected after cross-linking with BMB or BMH, but not with BMOE. **f**, Structures and estimated cross-linking distances for cross-linkers used in this study. dBBr: dibromobimane, BMOE: bismaleimidoethane, BMB: 1,4-bismaleimidobutane, BMH: bismaleimidohexane.

This way we generated FtsA double cysteine mutant strains *ftsA^3x HA, P98C, S118C^* and *ftsA^3x HA, E199C, S252C^*, which probe the lateral FtsA_i_-FtsA_i*−1_ and the longitudinal FtsA_i_-FtsA_i±1_ interface of the FtsA double filament, respectively (Figure 5b). *E. coli* cells were treated with bismaleimidoethane (BMOE) in exponential phase (OD_600_ = 0.2-0.4). BMOE enters living *E. coli* cells and rapidly cross-links closely spaced thiols such as cysteine side chains *in vivo*. FtsA species were visualised by SDS-PAGE and Western blotting against the 3x HA-tag in FtsA (Figure 5c). We detected efficient formation of crosslinked FtsA dimers for both the *ftsA^3x HA, P98C, S118C^* and *ftsA^3x HA, E199C, S252C^* double mutants, but only weak background signal spread across multiple species in single cysteine mutant controls. We also detected higher order polymers for the *ftsA^3x HA, E199C, S252C^* mutant since the open symmetry of the longitudinal filament contact allows for more than two FtsA monomers to be cross-linked, through chaining.

To probe the lateral association *in vivo* further, we generated the two, additional FtsA single cysteine mutant strains *ftsA^3x HA, D123C^* and *ftsA^3x HA, Q155C^*, which probe the lateral FtsA_i_-FtsA_i*_ and FtsA_i_-FtsA_i*−1_ interface of the FtsA double filament, respectively (Figure 5d). These single cysteine mutants may cross-link to their symmetry mates because of the local C2 symmetry in each of the two lateral filament interfaces. However, C_β_-C_β_ distances in the VmFtsA double filament structure of 3.9 Å for *ftsA^3x HA, D123C^* and 12.6 Å for *ftsA^3x HA, Q155C^* indicated that cross-linking with BMOE with an expected cross-linking distance of ∼8 Å might be inefficient, which prompted us to try thiol-directed cross-linkers of different lengths (Figure 5f). The *ftsA^3x HA, D123C^* mutant showed more efficient cross-linking of FtsA using dibromobimane (dBBr) than using BMOE (Figure 5e, left). In case of the *ftsA^3x HA, Q155C^* mutant, BMOE cross-linking did not lead to efficient formation of covalent FtsA dimers, whereas treatment with the longer maleimide cross-linkers 1,4-bismaleimidobutane (BMB) and bismaleimido-hexane (BMH) did (Figure 5e, right). Taken together, our data strongly suggest that FtsA forms protofilaments in cells and, that these protofilaments are further arranged into antiparallel double filaments as suggested by the VmFtsA^1-396^ crystal structure (PDB 7Q6F).

## DISCUSSION

Here, we found that FtsA forms antiparallel double filaments in *E. coli* and that *in vitro* the same filaments are induced by the binding of the cytoplasmic tail of FtsN. The only other polymerising actin-like protein known to form antiparallel and, therefore, apolar double filaments is the actin homologue of the elongasome, MreB, which serves as a “rudder” (Hussain *et al.*, 2018; van den Ent *et al.*, 2014; Wagstaff and Löwe, 2018). Given the shared preference for negative Gaussian curvature displayed by FtsA-FtsN and MreB double filaments reconstituted in and on liposomes (Hussain *et al.*, 2018; Salje *et al.*, 2011; van den Ent *et al.*, 2014) (Figure 4), we propose that MreB and FtsA share the same curvature sensing mechanism. Hence, we put forward a model for curvature-guided cell constriction by FtsA-FtsN double filaments, which serve as a “rudder” to align the direction of the divisome’s PG synthesis activity, more precisely glycan strand synthesis, with the cell’s circumference (Figure 6a).

**Figure 6.**
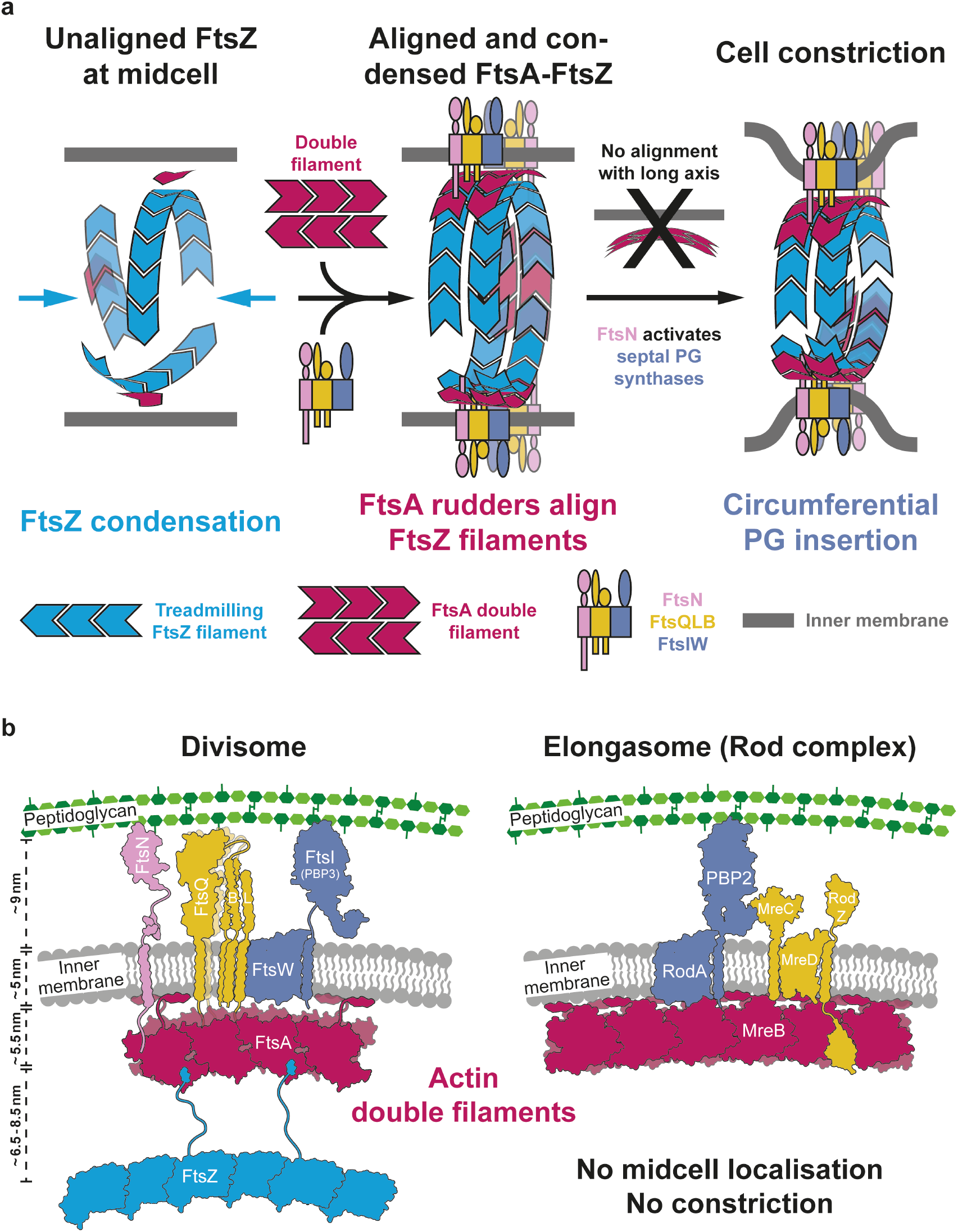
Model for curvature-guided septal PG synthesis and cell constriction mediated by FtsA double filaments. **a**, Logical steps towards a divisome primed for curvature-guided septal PG synthesis. The temporal order of events remains to be determined. FtsZ filaments recruited to midcell are unaligned (left). Downstream divisome proteins are recruited and condensed into the narrow midcell plane by treadmilling FtsZ filaments that themselves will also be aligned through their interaction with curvature-sensing FtsA filaments (middle). Condensed complexes are aligned with the short axis of the cell by FtsA double filaments because of their curvature-sensing mechanism, which we propose here they share with MreB. Finally, FtsZ filaments distribute divisome components via treadmilling and might thereby reinforce their alignment with the short axis of the cell. FtsN is required for FtsA double filament formation and activation of septal PG synthases. By coordinating both activities across the inner membrane, FtsN might constitute a synchronising activation switch for the divisome, which allows circumferential synthesis of septal peptidoglycan through the alignment activity, and at the same time cell constriction to commence through the activation of PG synthesis (right). **b**, Schematic overview of core components of the divisome and elongasome highlighting the evolutionary relationship between the two complexes. We propose here that both complexes utilise the curvature-sensing properties of their cytoplasmic actin double filament scaffolds (red) to direct peptidoglycan synthesis around the cell’s circumference. The bipartite PG synthases (FtsWI and RodA-PBP2, purple) are connected to the actin double filaments via integral membrane proteins that serve as structural and regulatory subunits (yellow), or potentially even directly. Unlike the elongasome, the divisome, in FtsN, possesses an additional regulatory subunit that might be necessary since cell division is regulated during the cell cycle. FtsZ is absent in the elongasome. FtsZ localises the divisome and its activities to midcell and probably more importantly into a narrow plane, an activity that is neither required nor desired in the elongasome.

In our model, we discriminate three distinct phases in divisome assembly and maturation: unaligned FtsZ filaments at midcell, a fully assembled divisome aligned with the short axis of the cell, and a fully activated divisome synthesising septal PG and driving cell constriction (Figure 6a). However, the temporal order of individual recruitment and activation events during divisome maturation remains largely unknown and would benefit from *in vivo* single molecule imaging of FtsA. FtsZ filaments are positioned at midcell to determine the division plane but are unaligned in the absence of an alignment mechanism (Figure 6a, left). After recruitment of divisome proteins and Z-ring condensation (Squyres *et al.*, 2021; Whitley *et al.*, 2021), FtsN-induced FtsA double filaments will align themselves and the other divisome components with the short axis of the cell, possibly aided by FtsZ treadmilling, which may provide a distribution mechanism that avoids FtsA filaments and divisome components to cluster in one spot (Bisson-Filho *et al.*, 2017; McQuillen and Xiao, 2020; Yang *et al.*, 2017) (Figure 6a, middle). We propose that curvature-sensing FtsA double filaments provide a solution to the FtsZ alignment problem noted previously (Du and Lutkenhaus, 2019). Consistently, the fraction of directionally treadmilling FtsZ filaments was reported to be decreased in a Δ*ftsA* strain of *B. subtilis* (Squyres *et al.*, 2021). Together FtsA double filaments and treadmilling FtsZ filaments we suggest align and evenly distribute divisome components in the narrow division plane. Most importantly, this will restrict movement of FtsWI, the bipartite PG synthase of the divisome, in such a way that cell-constricting septal PG synthesis follows the cell’s circumference (Figure 6a, right). In this context, FtsN might function as an activation switch of the divisome by coordinating activities of FtsA and FtsWI. Recently, a more direct interaction between FtsA and FtsW has been proposed (Park *et al.*, 2021).

Based on bacterial and yeast two-hybrid assays, it has been suggested that FtsN promotes divisome maturation through depolymerisation of FtsA, which would free FtsA’s IC domain and thereby allow recruitment of downstream divisome components (Pichoff *et al.*, 2012; Pichoff *et al.*, 2015, 2018). In contradiction to that we found that the short cytoplasmic tail of FtsN promotes FtsA polymerisation, or more precisely FtsA double filament formation (Figure 2c, d and g). Our work adds to previous evidence illustrating that FtsA can form different polymers (Krupka *et al.*, 2017; Schoenemann *et al.*, 2018; Szwedziak *et al.*, 2012). Using FRET probes attached to FtsA, a complementary fluorescence microscopy study found FtsA* to be less polymeric than wildtype FtsA on supported lipid bilayers (Radler *et al.*, manuscript in preparation). Addition of cytoplasmic FtsN peptide however induced FtsA* polymerisation to levels comparable to wildtype FtsA, again questioning the prevalent model that FtsN depolymerises FtsA. Further, our data clearly demonstrate that the FtsA double filament is compatible with FtsN binding, as could be the case for other downstream divisome components. Double filament formation will also position FtsA’s IC domain close to the inner membrane (Supplementary Figure S1c), which could facilitate recruitment of downstream divisome components that possibly bind to FtsA’s IC domain such as FtsQ (Baranova *et al.*, 2020).

Taken together with recent reports establishing FtsWI (Taguchi *et al.*, 2019) and RodA-PBP2 (Cho *et al.*, 2016; Sjodt *et al.*, 2020) as bipartite PG synthases, our data on the similarities between FtsA and MreB double filaments strengthen the previously proposed evolutionary relationship between the divisome and elongasome (Szwedziak and Löwe, 2013) (Figure 6b). In Chlamydia, one of the few bacteria that lack FtsZ, MreB was implied to organise cell division (Jacquier *et al.*, 2015; Ranjit *et al.*, 2020), yet another indication that divisome and elongasome share at least some basic principles of function.

In this context it is also interesting to speculate which of FtsZ’s many functions in cell division “converted” the presumably ancestral elongasome into the divisome. Our model suggests that FtsZ is the long-range organiser of the division site which ensures that septal PG synthases are functioning in a single division plane, only, and are evenly distributed around the cell’s circumference. FtsA on the other hand aligns these PG synthases with the orientation of the division ring, making sure septal PG glycan synthesis by the divisome goes around the ring’s circumference. It will require further studies before we truly understand, or can reconstitute FtsZ-based cell division but our present study highlights the central role FtsA polymerisation might play in this process and, further, illustrates important similarities between elongasome and divisome mechanisms.

## METHODS

### Expression plasmids

Expression plasmids (Supplementary Table T 2) were cloned by isothermal assembly using NEBuilder HiFi DNA Assembly Mix (NEB). Plasmids were transformed into *E. coli* MAX Efficiency DH5α (ThermoFisher) or C41(DE3) cells (Lucigen or Sigma) for plasmid propagation and protein expression, respectively.

### Protein expression and purification

The amino acid sequences of all proteins used in this study are listed in Supplementary Table T 3. If applicable, removable tags that were cleaved during purification are indicated. All purification steps were carried out on ice or in a cold room at 4-6°C unless stated otherwise. Buffers were prepared in Millipore water (MPW), pH-adjusted at room temperature and filtered through a 0.2 μm PES filter.

#### SUMO protease (GST-SENP1)

GST-SENP1 (pTN_AN_902) was expressed in C41(DE3) cells. Cells were grown in 2xTY medium, supplemented with 100 µg/ml ampicillin, at 37°C. Expression was induced at OD_600_ = 0.6-0.8 by adding 0.5 mM IPTG. Cells were grown over night at 18°C and harvested by centrifugation. Cells were lysed in buffer SA (50 mM Tris/HCl, 150 mM NaCl, 2 mM TCEP, 1 mM EDTA, 5 % glycerol, pH 8.5), supplemented with DNase, RNase and cOmplete EDTA-free Protease Inhibitor Cocktail (Roche), using a cell disruptor at 25 kpsi (Constant Systems). The lysate was cleared by ultracentrifugation at 100,000x g for 30 min at 4°C. The supernatant was added to pre-washed Glutathione Sepharose 4B beads (Cytiva) and incubated for 2h at 4°C, with gentle stirring. The sample was passed through an Econo gravity flow column (Bio-Rad) and washed 5x with 50 ml buffer SA, followed by 2x 50 ml buffer SB (buffer SA with 500 mM NaCl) and 2x 50 ml buffer SA. The protein was eluted in 5x 5 ml buffer SA supplemented with 10 mM reduced glutathione. Peak fractions were pooled and concentrated with a Vivaspin 20 concentrator (30 kDa MWCO, Sartorius). The protein was further purified by size exclusion chromatography on a HiLoad 26/600 Superdex 200 pg column (Cytiva), equilibrated in buffer SEC-S (50 mM Tris/HCl, 50 mM NaCl, 5 mM TCEP, 1 mM EDTA, 1 mM NaN_3_, 5 % glycerol, pH 8.0). Peak fractions were analysed by SDS-PAGE, pooled and concentrated to ∼15-20 mg/ml using Vivaspin 20 concentrators (30 kDa MWCO, Sartorius). Small aliquots were flash-frozen in liquid nitrogen and stored at −80°C.

#### TEV protease (6H-PolG*)

6H-PolG* (pTN_AN_901) was expressed in C41(DE3) cells. Cells were grown in 2xTY medium, supplemented with 30 µg/ml kanamycin, at 37°C. Expression was induced at OD_600_ = 0.6-0.8 by adding 1 mM IPTG. Cells were grown over night at 19°C and harvested by centrifugation. Cells were lysed in TEV lysis buffer (20 mM Tris/HCl, 500 mM NaCl, 1 mM TCEP, 1 mM NaN_3_, pH 8.0), supplemented with DNase and RNase using a cell disruptor at 25 kpsi (Constant Systems). The lysate was cleared by ultracentrifugation at 100,000x g for 30 min at 4°C. 20 mM imidazole was added to the supernatant before loading onto a HisTrap FF column (Cytiva). The column was washed with TEV lysis buffer supplemented with 20 mM imidazole. The protease was eluted in TEV lysis buffer with increasing amounts of imidazole (50/100/200/500 mM steps). Peak fractions were pooled and concentrated with Vivaspin 20 concentrators (10 kDa MWCO, Sartorius). Concentrated protein was further purified by size exclusion chromatography on a HiLoad 16/600 Superdex 75 pg column (Cytiva), equilibrated in TEV lysis buffer. Peak fractions were analysed by SDS-PAGE, pooled and concentrated to 30 mg/ml using Vivaspin 20 concentrators (10 kDa MWCO, Sartorius). Small aliquots were flash-frozen in liquid nitrogen and stored at −80°C.

#### Full-length FtsAs

C-terminal intein-CBD-12H fusions of EcFtsA (pTN_AN_001), EcFtsA^M96E, R153D^ (pTN_AN_059), EcFtsA^E124A^ (pTN_AN_003), EcFtsA^I143L^ (pTN_AN_004), EcFtsA^G50E^ (pTN_AN_024), EcFtsA^R286W^/EcFtsA* (pTN_AN_005) or VmFtsA (pTN_AN_057) were expressed in C41(DE3) cells. Cells were grown in 2xTY, supplemented with 100 µg/ml ampicillin, at 37°C. Expression was induced at OD_600_ = 0.8-1.0 by adding 0.5 mM IPTG. Cells were grown over night at 18°C and harvested by centrifugation. Cells were lysed in buffer LB2 (50 mM Tris/HCl, 500 mM NaCl, 5 mM TCEP, 10 mM MgCl2, 1 mM NaN_3_, pH 8.0), supplemented with DNase, RNase and cOmplete EDTA-free Protease Inhibitor Cocktail (Roche), using a cell disruptor at 25 kpsi (Constant Systems). The lysate was cleared by ultracentrifugation at 100,000x g for 30 min at 4°C. 50 mM imidazole was added to the supernatant before loading onto a HisTrap HP column (Cytiva). After loading, the column was washed with buffer SEC2 (50 mM CHES/KOH, 500 mM KCl, 5 mM TCEP, 10 mM MgCl2, 5 % glycerol, 1 mM NaN_3_, pH 9.0) supplemented with 100 mM imidazole. The protein was eluted in buffer SEC2 with increasing amounts of imidazole (200/300/500 mM steps). Most of the protein eluted in 200 mM imidazole. Peak fractions were pooled and loaded onto chitin resin (NEB) packed in a XK 50/20 column (Cytiva). The column was thoroughly washed with buffer CHIT2 (50 mM CHES/KOH, 500 mM KCl, 5 mM TCEP, 10 mM MgCl2, 5 % glycerol, 1 mM EGTA, 1 mM NaN_3_, pH 9.0), followed by 2 column volumes (CV) of buffer CHIT2 + 50 mM β-mercaptoethanol. Afterwards, the flow was stopped, and the column incubated over night at 4°C, allowing for intein cleavage. Cleaved FtsA was eluted with 2 CV of buffer CHIT2 + 50 mM β-mercaptoethanol, followed by 2 CV of buffer CHIT2. The protein was concentrated with Vivaspin 20 concentrators (30 kDa MWCO, Sartorius). Concentrated protein was further purified by size exclusion chromatography on a HiLoad 16/600 Superdex 200 pg column (Cytiva), equilibrated in buffer SEC2. Peak fractions were analysed by SDS-PAGE, pooled and concentrated to 7-8 mg/ml using Vivaspin 20 concentrators (30 kDa MWCO, Sartorius). Small aliquots were flash-frozen in liquid nitrogen and stored at −80°C. Protein mass for each batch was verified by ESI-TOF mass spectrometry on a Micromass LCT mass spectrometer (Waters).

#### C-terminally truncated FtsAs

N-terminal 6H-SUMO fusions of EcFtsA^1-405^ (pTN_AN_022), XpFtsA^1-396^ (pTN_AN_050) or VmFtsA^1-396^ (pTN_AN_052) were expressed in C41(DE3) cells. Cells were grown in 2xTY, supplemented with 30 µg/ml kanamycin, at 37°C. Expression was induced at OD_600_ = 0.8-1.0 by adding 0.5 mM IPTG. For XpFtsA_ΔC_, cells were harvested by centrifugation 5h later. Cells expressing 6H-SUMO-EcFtsA^1-405^ or 6H-SUMO-VmFtsA^1-396^ were grown over night at 25°C and harvested the next day by centrifugation. Cells were lysed in buffer LB2 (50 mM Tris/HCl, 500 mM NaCl, 5 mM TCEP, 10 mM MgCl_2_, 1 mM NaN_3_, pH 8.0), supplemented with DNase, RNase and cOmplete EDTA-free Protease Inhibitor Cocktail (Roche), using a cell disruptor at 25 kpsi (Constant Systems). The lysate was cleared by ultracentrifugation at 100,000x g for 30 min at 4°C. 20 mM imidazole was added to the supernatant before loading onto a HisTrap HP column (Cytiva). After loading, the column was washed with buffer SEC2 (50 mM CHES/KOH, 500 mM KCl, 5 mM TCEP, 10 mM MgCl_2_, 5 % glycerol, 1 mM NaN_3_, pH 9.0) supplemented with 20 mM imidazole. The protein was eluted in buffer SEC2 with increasing amounts of imidazole (100/300/500 mM steps). Most of the protein eluted in 100 mM imidazole. Peak fractions were pooled and mixed with pre-washed Glutathione Sepharose 4B beads (Cytiva) and purified GST-SENP1 (final concentration ∼0.01 mg/ml). The sample was incubated over night at 4°C, with gentle stirring. The next day, the sample was passed through an Econo gravity flow column (Bio-Rad). The flow-through was concentrated using Vivaspin 20 concentrators (30 kDa MWCO, Sartorius). Concentrated protein was further purified by size exclusion chromatography on a HiLoad 26/600 Superdex 200 pg column (Cytiva), equilibrated in buffer SEC2 in case of EcFtsA^1-405^ or crystallisation buffer (20 mM CHES/KOH, 100 mM KCl, 5 mM TCEP, 5 mM MgCl_2_, 5 % glycerol, 1 mM NaN_3_, pH 9.0) in case of XpFtsA^1-396^ and VmFtsA^1-396^. Peak fractions were analysed by SDS-PAGE, pooled and concentrated to 12-19 mg/ml using Vivaspin 20 concentrators (30 kDa MWCO, Sartorius). Small aliquots were flash-frozen in liquid nitrogen and stored at −80°C. Protein mass for each batch was verified by ESI-TOF mass spectrometry on a Micromass LCT mass spectrometer (Waters).

#### Isotope-labelled VmFtsN^1-29^-ENLYFQ and Gly-Gly-VmFtsN^2-29^

Isotope-labelled proteins were overexpressed in C41(DE3) cells in M9 media (6 g/L Na_2_HPO_4_, 3 g/L KH_2_PO_4_, 0.5 g/L NaCl) supplemented with 1.7 g/L yeast nitrogen base without NH_4_Cl and amino acids (Sigma Y1251). 1 g/L ^15^NH_4_Cl and 4 g/L ^13^C-glucose were supplemented for ^15^N and ^13^C labelling, respectively. VmFtsN^1-29^-ENLYFQ (pTN_AN_069) and Gly-Gly-VmFtsN^2-29^ (pTN_AN_071) were expressed in C41(DE3) cells grown in the presence of 30 µg/ml ampicillin at 37°C. Expression was induced at OD600 = 0.6-0.8 by adding 0.5 mM IPTG. Cells were grown over night at 25°C and harvested by centrifugation. Cells were lysed in buffer LB5 (50 mM HEPES/KOH, 250 mM KCl, 1 mM TCEP, pH 7.7), supplemented with DNase, RNase and cOmplete EDTA-free Protease Inhibitor Cocktail (Roche), using a cell disruptor at 25 kpsi (Constant Systems). The lysate was cleared by ultracentrifugation at 100,000x g for 30 min at 4°C. 20 mM imidazole was added to the supernatant before loading onto a HisTrap HP column (Cytiva). The column was washed with buffer LB5 supplemented with 20 mM or 50 mM imidazole for VmFtsN^1-29^-ENLYFQ and Gly-Gly-VmFtsN^2-29^, respectively. Proteins were eluted in buffer LB5 with increasing amounts of imidazole (100/200/300/500 mM steps). Peak fractions were pooled and mixed with TEV protease, 6H-PolG*, (final concentration 0.01-0.02 mg/ml). The sample was incubated over night at 4°C, with gentle stirring. The next day, samples were diluted with buffer LB5 to a final imidazole concentration of ∼50 mM and loaded onto a HisTrap HP column equilibrated in buffer LB5 supplemented with 50 mM imidazole. The column was washed with buffer LB5 supplemented with 50 mM imidazole. Flow-through and wash fractions were pooled and concentrated using Vivaspin 15R concentrators (2 kDa MWCO HY, Sartorius).

For Gly-Gly-VmFtsN^2-29^ only, an additional cation exchange purification step was performed. The sample was diluted in buffer SA2 (50 mM HEPES/KOH, 50 mM KCl, 1 mM TCEP, pH 7.7) to a final salt concentration of ≤ 100 mM and loaded on a HiTrap SP HP column (Cytiva). The sample was eluted with a linear gradient of buffer SB2 (50 mM HEPES/KOH, 1000 mM KCl, 1 mM TCEP, pH 7.7). Peak fractions were pooled and concentrated.

Concentrated VmFtsN^1-29^-ENLYFQ or Gly-Gly-VmFtsN^2-29^ was further purified by size exclusion chromatography on a Superdex Peptide 10/300 GL column (Cytiva), equilibrated in NMR buffer (50 mM MES/KOH, 50 mM KCl, 1 mM TCEP, pH 6.0). Peak fractions were analysed by SDS-PAGE, pooled and concentrated to 2-3 mg/ml using Vivaspin 15R concentrators (2 kDa MWCO HY, Sartorius). Small aliquots were flash-frozen in liquid nitrogen and stored at −80°C. Protein mass for each batch was verified by ESI-TOF mass spectrometry on a Micromass LCT mass spectrometer (Waters).

#### FtsN peptides

All FtsN peptides were chemically synthesised by Generon/Neobiotech. Purity (≥ 95 % in HPLC) and molecular mass were verified by the company. Lyophilised peptides were resuspended in binding buffer (50 mM HEPES/KOH, 100 mM KAc, 5 mM MgAc_2_, pH 7.7). Stock concentrations were determined using an ND-1000 spectrophotometer (NanoDrop Technologies) or, if the peptide did not contain any tyrosines or tryptophans, a Direct Detect Infrared Spectrometer (Merck Millipore). Concentrations of resuspended peptides were in the range of 20-70 mM. Peptide sequences are given in Supplementary Table T 4.

### Crystal structure determination

Crystallisation conditions were screened using our in-house high-throughput crystallisation facility (Gorrec and Löwe, 2018). EcFtsA^1-405^ at 7 mg/ml, XpFtsA^1-396^ at 7 mg/ml or VmFtsA^1-396^ at 5 mg/ml was mixed with 2 mM ATP. For co-crystallisation, VmFtsA^1-396^ at 3 mg/ml was mixed with 2 mM ATP and 0.353 mM VmFtsN^1-29^ (5-fold molar excess). For EcFtsA^1-405^ and XpFtsA^1-396^, 100 nl protein solution and 100 nl of the crystallisation solutions were mixed in MRC sitting drop crystallisation plates for vapour diffusion. 500 nl protein solution and 500 nl of crystallisation solutions were used to obtain optimised co-crystals of VmFtsA^1-396^ with VmFtsN^1-29^. Optimisation for VmFtsA^1-396^ followed the idea of an “anticipated optimisation approach” (Gorrec, 2019). 250 nl of protein solution were mixed with 200 nl of initial crystallisation solution (31.55 % v/v PEG 400, 0.21 M MgCl_2_, 0.1 M Tris/HCl pH 8.5) and 50 nl of follow-up crystallisation solution (10 % v/v 2-propanol, 0.2 M CaAc_2_, 0.1 M MES/NaOH pH 6.0). When completed, drops were pipetted up and down 5 times by the Mosquito robot (TTP Labtech) to minimise diffusion effects. Plates were incubated at 21°C. 1-2 µl of cryoprotectant solution were added to crystals shortly before mounting. Single crystals were mounted in CryoLoops (Hampton Research) and flash-frozen in liquid nitrogen. VmFtsA^1-396^-FtsN^1-29^ co-crystals (PDB 7Q6I) were soaked in cryoprotectant solution for 1h prior to mounting. Optimised conditions yielding crystals and cryoprotectant solutions are listed in Supplementary Table T 1. Diffraction data were collected on single crystals at Diamond Light Source (DLS), Harwell, UK, at 100 K, as indicated in Supplementary Table T 1. XDS (Kabsch, 2010), POINTLESS and SCALA (Evans, 2006) or DIALS (Winter *et al.*, 2018) were used to process diffraction images. Initial phases were obtained by molecular replacement using PHASER (McCoy *et al.*, 2007). Search models used for molecular replacement are indicated in Supplementary Table T 1. Models were rebuilt manually using MAIN (Turk, 2013) and COOT (Emsley *et al.*, 2010) and refined using REFMAC (Murshudov *et al.*, 1997) and PHENIX (Afonine *et al.*, 2018). Models were validated using PROCHECK (Laskowski *et al.*, 1993) and MOLPROBITY (Williams *et al.*, 2018). Final statistics are summarised in Supplementary Table T 1, and the structure factors as well as atomic coordinates have been deposited in the Protein Data Bank (PDB) with accession codes 7Q6D, 7Q6G, 7Q6F and 7Q6I. Note that VmFtsN^1-29^ (chains X and Y) density was modelled as VmFtsN^1-8^ for refinement but deposited as polyAla.

### Surface plasmon resonance (SPR)

SPR was performed using a Biacore T200 using CM5-sensor chips (Cytiva). Both reference control and analyte channels were equilibrated in binding buffer (50 mM HEPES/KOH, 100 mM KAc, 5 mM MgAc_2_, pH 7.7). EcFtsA, EcFtsA^1-405^ or VmFtsA^1-396^ was immobilised onto the chip surface via amide coupling using the supplied kit (Cytiva) to reach a RU value of between 2000 and 7800 RU for separate experiments. SPR runs were performed with analytes injected for 120 s followed by a 300 s dissociation in a 1:2 dilution series with initial concentrations of: 10 µM for peptide EcFtsN^1-32^ (Figure 2a); 60 µM for VmFtsN^1-29^ (Supplementary Figure S2a); 20 µM for peptides EcFtsN^1-12^, EcFtsN^1-22^, EcFtsN^11-32^, EcFtsN^4-26^, EcFtsN^1-32^, EcFtsN^1-32, D5N^, EcFtsN^1-32, ΔRK1^, EcFtsN^1-32, ΔRK2^, EcFtsN^1-32, ΔRK3^, EcFtsN^1-32, ΔRK2,3^ and EcFtsN^1-33, scrambled^ (Supplementary Figure S2i). After reference and buffer signal correction, sensogram data were fitted using Prism 8.0 (GraphPad Software Inc). The equilibrium response (*R_eq_*) data were fitted to a single-site interaction model to determine *K_d_*:

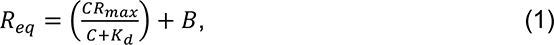

where *C* is the analyte concentration and *R_max_* is the maximum response at saturation and *B* is the background resonance. Binding constants are given as mean.

### Fluorescence polarisation (FP)

Peptides EcFtsN^1-32^-C and VmFtsN^1-29^-C were labelled with maleimide-Atto 495 (Merck Life Science UK). EcFtsA^1-405^ and VmFtsA^1-396^ were buffer exchanged into binding buffer (50 mM HEPES/KOH, 100 mM KAc, 5 mM MgAc_2_, pH 7.7) using Zeba Spin Desalting Columns (0.5 ml, 7 kDa MWCO, ThermoFisher) prior to FP experiments. 20 nM labelled peptide was mixed 1:1 with a 1:2 dilution series of protein with initial concentrations of 40 µM for EcFtsA^1-405^ or 120 µM for VmFtsA^1-396^ in binding buffer supplemented with Tween-20 (50 mM HEPES/KOH, 100 mM KAc, 5 mM MgAc_2_, pH 7.7, 0.05 % v/v Tween-20) in a 384-well low flange black flat bottom non-binding surface microplate (Corning). Reactions were prepared in triplicate. Fluorescence polarisation was measured using a PHERAstar FSX (BMG Labtech) directly after initial reaction setup and after incubation at room temperature for 30 min and 2 h to ensure equilibrium had been reached. Error bars indicate the standard deviation of three technical replicates. Data were fitted using Prism 8.0 (GraphPad Software Inc). Dissociation constants were calculated using a two-step model:

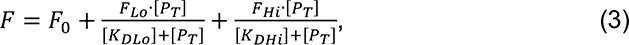

where *F_Lo_* and *F_Hi_* are the anisotropy changes at saturation of low and high affinity sites with binding constants of *K_DLo_* and *K_DHi_*, respectively. Binding constants were averaged from 5 replicates and are given as mean ± SEM.

### Fluorescence detection system-analytical ultracentrifugation (FDS-AUC)

Peptides EcFtsN^1-32^-C and VmFtsN^1-29^-C were labelled with maleimide-Atto 495 (Merck Life Science UK). EcFtsA^1-405^ and VmFtsA^1-396^ were buffer exchanged into binding buffer (50 mM HEPES/KOH, 100 mM KAc, 5 mM MgAc_2_, pH 7.7) using Zeba Spin Desalting Columns (0.5 ml, 7 kDa MWCO, ThermoFisher) prior to FDS-AUC experiments. 20 nM labelled peptide was mixed 1:1 with a 1:3 dilution series of protein with initial concentrations of 90 µM for EcFtsA^1-405^ or 120 µM for VmFtsA^1-396^ in binding buffer supplemented with Tween (50 mM HEPES/KOH, 100 mM KAc, 5 mM MgAc_2_, pH 7.7, 0.05 % v/v Tween-20). Samples were centrifuged at 45,000 rpm at 20°C in an An50Ti rotor using an Optima XL-I analytical ultracentrifuge (Beckman) equipped with a fluorescence optical system (Aviv Biomedical) with fixed excitation at 488 nm and fluorescence detection at > 505 nm. Data were processed and analysed using SEDFIT 16 and SEDPHAT 15.2 (Schuck, 2003) according to the published protocol for high-affinity interactions detected by fluorescence (Chaturvedi *et al.*, 2017). Binding constants were estimated using a two-step model. Data were plotted using GUSSI (Brautigam, 2015).

### Co-pelleting assay

EcFtsA^1-405^ or VmFtsA^1-396^ at 1 mg/ml (∼23 µM) alone or mixed with half, 1-, 3-, 6- or 10-fold the concentration of EcFtsN^1-32^ or VmFtsN^1-29^, respectively, in binding buffer (50 mM HEPES/KOH, 100 mM KAc, 5 mM MgAc_2_, pH 7.7) was incubated at room temperature for 5-15 min before centrifugation at 20,000x g for 10 min at 20°C. Supernatant and pellet fractions were separated carefully and analysed by SDS-PAGE. Protein band intensities were quantified using ImageJ 2.1.0 (Abràmoff *et al.*, 2004). Given are mean ± sd of two technical replicates for EcFtsA^1-405^ and VmFtsA^1-396^, respectively.

### Lipid monolayer assays and negative stain electron microscopy

Two-dimensional lipid monolayers were prepared from *E. coli* polar lipid extract (Avanti Polar Lipids) (Ford *et al.*, 2001; Levy *et al.*, 1999). Wells of a custom-made Teflon block were filled with 60 µl binding buffer (50 mM HEPES/KOH, 100 mM KAc, 5 mM MgAc_2_, pH 7.7). 20 µg of lipids, dissolved in chloroform, were applied on top of the buffer, and incubated for 2-3 min to allow evaporation of chloroform. Next, baked (60°C, overnight) CF300-Cu-UL EM grids (Electron Microscopy Sciences) were placed on top of the wells with the carbon side facing downwards. Grids were incubated for 20-60 min to allow formation and attachment of the lipid monolayer to the grid. In the meantime, full-length FtsA protein at 0.1 mg/ml (∼2.2 µM) for EcFtsA, EcFtsA^E124A^, EcFtsA^I143L^, EcFtsA^G50E^ and EcFtsA^R286W^/EcFtsA* or at 0.2 mg/ml (∼4.4 µM) for EcFtsA^M96E, R153D^ and VmFtsA was mixed with 1 mM ATP and indicated FtsN peptides at 10-fold molar excess, if not stated otherwise, in binding buffer (50 mM HEPES/KOH, 100 mM KAc, 5 mM MgAc_2_, pH 7.7). Samples were incubated for 10 min at RT. EM grids were carefully lifted off the buffer and blotted from the side using Whatman No1 filter paper. 4 µl sample were applied to the carbon side of an EM grid and incubated for 30 s before staining with 2% w/v uranyl formate. Grids were imaged on a Tecnai Spirit electron microscope (ThermoFisher) operating at 120 kV and equipped with a Gatan Orius SC200W camera. Presented micrographs were contrast adjusted and blurred for display purposes.

### Cryo-EM of EcFtsA-FtsN^1-32^ double filaments on lipid monolayer

Lipid monolayers were prepared as described above, with the exception that Quantifoil Au R0.6/1 300 mesh grids (Quantifoil) were used. EcFtsA at 0.1 mg/ml (∼2.2 µM) was mixed with 1 mM ATP and EcFtsN^1-32^ at 22 µM in binding buffer (50 mM HEPES/KOH, 100 mM KAc, 5 mM MgAc_2_, pH 7.7), and incubated for 20 min at RT. Grids were blotted from the side after attachment of the monolayer using Whatman No 1 filter paper and inserted into a Vitrobot Mark III (ThermoFisher) set to 20°C and 100% humidity. 3 µl of mixed protein sample were applied to the carbon side of the grid, incubated for 30 s and blotted for 12-15 s (0.5 s drain time, −15 blot force) before plunge-freezing into liquid ethane maintained at −180°C using a cryostat (Russo *et al.*, 2016). Grids were imaged using a Tecnai G2 Polara (ThermoFisher) operating at 300 kV. Movies were collected on a Falcon III direct electron detector at a pixel size of 1.38 Å, −3.3 to −4 µm defocus and a total dose of 100 e^−^/Å^2^ using EPU (ThermoFisher) for automated acquisition. Data were processed using MotionCor2 (Zheng *et al.*, 2017), CTFFIND4 (Rohou and Grigorieff, 2015) and RELION3 (Zivanov *et al.*, 2018). A total of 104,660 particles were automatically picked and extracted, with the presented 2D class average corresponding to 14,602 particles (Figure 2e). Presented micrographs were contrast adjusted and blurred for display purposes.

### Multiple sequence alignment

Annotated FtsA (TIGR01174) and FtsN (TIGR02223) sequences from 2110 bacterial genomes were retrieved from the AnnoTree server (Haft *et al.*, 2001; Mendler *et al.*, 2019; Parks *et al.*, 2018). Genomes with multiple or short FtsA or FtsN sequences were removed. The remaining 1983 FtsN sequences were aligned using Clustal Omega (Sievers *et al.*, 2011). Only sequence stretches corresponding to FtsN’s cytoplasmic and transmembrane domain (FtsN^cyto-TM^) were used for analysis, corresponding to EcFtsN^1-54^ for example. Only FtsN^cyto-TM^ sequences with a length between 44 and 64 amino acids and a sequence identity to EcFtsN^cyto-TM^ ≥ 35 % were retained if the sequence identity of the corresponding FtsA sequence to *E. coli* FtsA also was ≥ 70 %. The remaining 245 FtsN^cyto-TM^ sequences were aligned using Muscle (Edgar, 2004). The sequence logo was created using Weblogo 3.7.4 (Crooks *et al.*, 2004).

### Hydrogen deuterium exchange mass spectrometry (HDX-MS)

VmFtsA^1-396^ at 10 µM was mixed with 2 mM ATP and, if indicated, VmFtsN^1-29^ at 3- (30 µM) or 10-fold (100 µM) molar excess in binding buffer (50 mM HEPES/KOH, 100 mM KAc, 5 mM MgAc_2_, pH 7.7). 5 µL of protein solution were added to 40 µL of D_2_O buffer at room temperature for 3, 30, 300 and 1800 seconds. Reactions were quenched and frozen until further processing. Quenched protein samples were rapidly thawed and subjected to proteolytic cleavage by pepsin followed by reversed phase HPLC separation. Briefly, the protein was passed through a 2.1 x 30 mm, 5 µm Enzymate BEH immobilised pepsin column (Waters, UK) at 200 µL/min for 2 min. Peptic peptides were trapped and desalted on a 2.1 x 5 mm C18 trap column (Acquity BEH C18 Van-guard pre-column, 1.7 µm, Waters, UK). Trapped peptides were subsequently eluted over 12 min using a 5-36% gradient of acetonitrile in 0.1% v/v formic acid at 40 µL/min. Peptides were separated on a 100 mm x 1 mm, 1.7 µm Acquity UPLC BEH C18 reverse phase column (Waters, UK). Peptides were detected on a SYNAPT G2-Si HDMS mass spectrometer (Waters, UK) acquiring over a *m/z* range of 300 to 2000, with standard electrospray ionisation (ESI) source and lock mass calibrated using [Glu1]-fibrino peptide B (50 fmol/µL). The mass spectrometer was operated at a source temperature of 80°C and a spray voltage of 2.6 kV. Spectra were collected in positive ion mode. Peptides were identified by MS^e^ (Silva *et al.*, 2005) using a 5-36% gradient of acetonitrile in 0.1% v/v formic acid over 12 min. The resulting MS data were analysed using Protein Lynx Global Server software (Waters, UK) with an MS tolerance of 5 ppm. Mass analysis of the peptide centroids was performed using DynamX software (Waters, UK). Only peptides with a score >6.4 were considered. The first round of analysis and identification was performed automatically by the DynamX software; however, all peptides (deuterated and non-deuterated) were manually verified at every time point for the correct charge state, presence of overlapping peptides, and correct retention time. Deuterium incorporation was not corrected for back-exchange and represents relative, rather than absolute changes in deuterium levels. Changes in H/D amide exchange in any peptide may be due to a single or multiple amides within that peptide. Time points were prepared in parallel and data for individual time points were acquired on the mass spectrometer on the same day.

### Nuclear magnetic resonance (NMR) spectroscopy

Backbone amide peaks of VmFtsN^1-29^-ENLYFQ were assigned using 167 µM ^15^N, ^13^C-labelled peptide at 278 K in NMR buffer (50 mM MES/KOH, 50 mM KCl, 1 mM TCEP, pH 6.0). Standard triple resonance spectra: HNCO, HN(CA)CO, HNCACB, and CBCA(CO)NH (Bruker) were collected with 20 % non-uniform sampling and processed with compressed sensing using the MddNMR package (Jaravine *et al.*, 2008). Backbone resonances were assigned using MARS (Jung and Zweckstetter, 2004). Topspin 3.6.0 (Bruker) was used for processing of 2D data and NMRFAM-Sparky 1.47 (Lee *et al.*, 2015) was used for spectra analysis. Assignment of VmFtsN^1-29^-ENLYFQ was transferred to Gly-Gly-VmFtsN^2-29^ using 295 µM ^15^N-labelled peptide at 278 K in NMR buffer. Gly1, Ala2 and Asn3, which were not observed for VmFtsN^1-29^-ENLYFQ, were newly assigned. The first glycine (G0) of Gly-Gly-VmFtsN^2-29^ was not observed.

For binding studies, ^1^H, ^15^N BEST-TROSY spectra were acquired at 278 K on 50 µM Gly-Gly-VmFtsN^2-29^ mixed with an equimolar concentration of VmFtsA^1-396^ in NMR buffer. As sensitivity was compromised by the formation of FtsA polymers upon binding of FtsN peptide, multiple short experiments were acquired and added up to define the ideal time window for data analysis. Each spectrum was acquired with 128 scans and a recycle delay of 400 ms, with a final spectral resolution of 4.7 Hz per points. Relative peak intensities were normalised to the C-terminal residue Arg29 of Gly-Gly-VmFtsN^2-29^ and analysed as I_bound_/I_free_, with I_bound_ and I_free_ being the peak intensities of Gly-Gly-VmFtsN^2-29^ with (bound) and without (free) VmFtsA^1-396^, respectively.

### EcFtsA-FtsN^1-32^ filaments on liposomes

Liposomes were prepared from *E. coli* polar lipid extract (Avanti Polar Lipids) by extrusion using a Mini Extruder fitted with a polycarbonate membrane with 1 µm pore size (Avanti Polar Lipids) in binding buffer without magnesium (50 mM HEPES/KOH, 100 mM KAc, pH 7.7). Preformed liposomes at 1 mg/ml were mixed with 0.5 mM MgATP and proteins at the following concentrations: no proteins (Figure 4a, left), FtsA at 20 µM and FtsN^1-32^ at 200 µM (Figure 4a, top right) and FtsA at 5 µM and FtsN^1-32^ at 50 µM (Figure 4a, bottom right). Samples were incubated at room temperature for 30 min without proteins or 10 min with proteins, respectively. 3 µl of sample were applied to a freshly glow-discharged Quantifoil Au R2/2 200 mesh grid (Quantifoil), blotted for 3.5-7.5 s (0.5 s drain time, −15 blot force) and plunge-frozen into liquid ethane maintained at −180°C using a cryostat (Russo *et al.*, 2016) using a Vitrobot Mark III (ThermoFisher) set to 20°C and 100% humidity. Grids were imaged on a Tecnai F20 (ThermoFisher) equipped with a Falcon II direct electron detector or a Glacios (ThermoFisher) equipped with a Falcon III detector. Microscopes were operated at 200 kV and cryogenic temperature. Presented micrographs were motion-corrected (if collected on the Glacios), contrast adjusted and blurred for display purposes.

### EcFtsA-FtsN^1-32^ filaments inside liposomes

EcFtsA-FtsN^1-32^ filaments were encapsulated into liposomes by dilution of CHAPS-solubilised *E. coli* total lipid extract (Avanti Polar Lipids) (Szwedziak *et al.*, 2014). 50 µl of preincubated FtsA at 20 µM mixed with 200 µM FtsN^1-32^ and 0.5 mM MgATP in binding buffer without magnesium (50 mM HEPES/KOH, 100 mM KAc, pH 7.7) were added to 50 µl of *E. coli* total lipid extract (Avanti Polar Lipids) solubilised at 10 mg/ml in binding buffer without magnesium supplemented with 20 mM CHAPS (50 mM HEPES/KOH, 100 mM KAc, pH 7.7, 20 mM CHAPS). The sample was incubated at room temperature for 35 min, before it was gradually diluted with 500 µl of binding buffer without magnesium supplemented with 0.5 mM MgATP (50 mM HEPES/KOH, 100 mM KAc, pH 7.7, 0.5 mM MgATP) within 20 min. 3 µl of sample were applied to a freshly glow-discharged Quantifoil Au R2/2 200 mesh grid (Quantifoil), blotted for 5.5-7.5 s (0.5 s drain time, −15 blot force) and plunge-frozen into liquid ethane maintained at −180°C using a cryostat (Russo *et al.*, 2016) using a Vitrobot Mark III (ThermoFisher) set to 20°C and 100% humidity. Grids were imaged on a Tecnai F20 (ThermoFisher) equipped with a Falcon II direct electron detector, operating at 200 kV and cryogenic temperature. Rarely, EcFtsA-FtsN^1-32^ filaments were observed inside liposomes when added to the outside of preformed liposomes (Figure 4d), most likely due to membrane rearrangements during handling. Here, preformed liposomes extruded to 1 µm at 1 mg/ml were mixed with 0.5 mM MgATP, FtsA at 2.5 µM and FtsN^1-32^ at 25 µM. Presented micrographs were contrast adjusted and blurred for display purposes. For 2D class averages (Figure 4e), movies were collected on a Glacios with a Falcon III at a pixel size of 1.99 Å, −2.5 to −4 µm defocus and a total dose of 56 e^−^/Å^2^ using SerialEM (Mastronarde, 2005). Data were processed using MotionCor2 (Zheng *et al.*, 2017), CTFFIND4.1 (Rohou and Grigorieff, 2015) and RELION-3.1 (Zivanov *et al.*, 2018). Presented images were upscaled and blurred for display purposes.

### Strain construction

The general cloning and recombination strategy is illustrated in Supplementary Figure S4a. The helper plasmid pKW20 (NCBI ID: MN927219.1) (Wang *et al.*, 2016) was used for genome engineering. pKW20 encodes for a constitutively expressed tracrRNA as well as Cas9 and *λ*-Red components under control of a L-arabinose-inducible promotor. Acceptor strain SFB123 was created by integrating a *pheS^T251A,A294G^-hygR* double selection cassette downstream of the *lpxC* gene using *λ*-Red recombineering (Yu *et al.*, 2000). *pheS^T251A,A294G^/pheS\** is a negative selection marker encoding a mutant phenylalanyl-tRNA synthetase that confers toxicity through misincorporation of 4-chloro-phenylalanine (4-CP) during translation (Miyazaki, 2015). We found a long (5 kB) homologous region upstream of FtsA to benefit recombination efficiency. SFB143 was used as donor strain during conjugation. SFB143 is an *E. coli* MDS42 *thi^-^* strain transformed with the non-transferrable conjugative plasmid pJF146 (NCBI ID: MK809154.1) (Fredens *et al.*, 2019). pJF146 bears an *apR* resistance marker and a truncated nick site of the origin of transfer. SFB143 cells were made chemically competent enabling parallelised transformation (Inoue *et al.*, 1990). Shuttle plasmid pFB483 was designed with a pMB1 origin of replication, a *pheS^T251A,A294G^-hygR* double selection cassette, a CRISPR array targeting *ftsW* and the region just upstream of *secM*, and a *ccdB* toxin gene (outsert) flanked by *Bsa*I acceptor sites via Golden Gate assembly (Engler *et al.*, 2008). CRISPR arrays were designed to mediate scarless excision. pFB483 was propagated in a ccdB survival strain.

Targeting constructs were split into 2-3 modules for insertion of single or double point mutations, respectively. Initially, the internal 3x HA-tag (120 bp including a *Xho*I restriction site) was inserted into *ftsA* using three modules, resulting in sTN001 (*ftsA269-3x HA-tag-270(SW), lpxC::neoR*), and propagated in modules in the following. Point mutations were introduced by PCR and modules assembled into pFB483 via Golden Gate assembly with *Bsa*I. The final targeting construct also introduced a kanamycin-selectable *neoR* marker downstream of the *lpxC* gene. Assembled shuttle vectors were transformed into SFB143 and selected on LB agar plates supplemented with 200 µg/ml hygromycin-B and 50 µg/ml apramycin at 37°C. Acceptor cells (SFB123) were grown to stationary phase in 5 ml LB supplemented with 10 µg/ml tetracycline. 4 ml of culture were harvested by centrifugation and washed three times in LB, before being transferred into 50 ml LB supplemented with 10 µg/ml tetracycline and 0.5 % w/v L-arabinose. Cells were grown at 37°C for 1h, harvested by centrifugation and washed three times in LB. In the meantime, donor transformants were washed off the agar plates using 2 ml LB and left at room temperature. All cultures were resuspended in LB to an OD_600_ of 40. 12.5 µl of acceptor cells were mixed 87.5 µl of donor cells and spotted onto well-dried TYE plates in 10-20 µl drops. Spots were air-dried, before plates were incubated at 30°C for 1h. Cells were washed off the plates with LB and transferred into 50 LB supplemented with 12.5 µg/ml kanamycin and 10 µg/ml tetracycline. Cells were grown at 37°C for 1h, harvested by centrifugation and plated on LB agar plates supplemented with 12.5 µg/ml kanamycin, 10 µg/ml tetracycline, 2 % glucose and 2.5 mM 4-CP. Strains were single colony purified, and verified by marker analysis and colony PCR followed by *Xho*I digestion and Sanger sequencing. Strains with desired point mutations were cured of pKW20 by repeated growth in LB in absence of antibiotics, diluted 1:10^6^ and plated on TYE plates. pKW20 was typically lost in one of eight cells after 2-3 growth cycles. Strains were verified by marker analysis and Sanger sequencing of PCR products covering the targeting region. Strains used in Figure 5c were further whole genome sequenced on a MiSeq (Illumina). NGS data were analysed using breseq v0.35.1 (Deatherage and Barrick, 2014).

Strains are listed in Supplementary Table T 5.

### Assessment of growth and cell elongation phenotypes

#### Growth on solid media

Strains were streaked on the same TYE plate and incubated at 37°C overnight. The next morning, strains were re-streaked on a fresh TYE plate and incubated at 37°C for 12h.

#### Growth in liquid media

Strains were inoculated into 5 ml LB and incubated at 37°C overnight. Cells were diluted 1/1000 into fresh LB into a 96-well flat bottom plate in octuplicate. The plate was incubated at 37°C in a Tecan microplate reader with regular shaking. Absorbance at 600 nm wavelength was measured every 5 min for 24h. OD_600_ values were background corrected, normalised to the maximum OD_600_ value of each well, averaged and plotted as mean ± sd.

#### DIC imaging of exponential phase cultures

Strains were inoculated into 5 ml LB and incubated at 37°C overnight. The next day, cells were diluted 1/1000 in 5 ml fresh medium and incubated at 37°C. 2-3 µl of exponential phase cells (OD_600_ = 0.2-0.3) were applied onto an agarose pad and imaged on a Nikon Eclipse E800 microscope equipped with a 100x oil objective and a Photometrics Iris 9 CMOS camera using a differential interference contrast (DIC) imaging setup. Presented images were contrast-adjusted for display purposes.

#### In vivo cysteine cross-linking

Strains were grown to exponential phase (OD_600_ = 0.2-0.4) and 0.9375 OD units harvested using centrifugation at 4°C. Cells were kept on ice for the duration of the experiment, unless stated otherwise. Cells were washed in 500 µl PBS, spun and resuspended in 50 µl PBS. 1.25 µl DMSO or cross-linker (dBBr: Dibromobimane, BMOE: Bismaleimidoethane, BMB: 1,4-bismaleimidobutane or BMH: Bismaleimido-hexane) in DMSO (20 mM stock) were added. Samples were incubated for 10 min and quenched by adding 1 µl 2-mercaptoethanol (2-ME, 1.43 M stock in MPW). Cells were spun and resuspended in 50 µl lysis buffer [1 mM EDTA (pH 7.4), 14.3 mM 2-ME, cOmplete EDTA-free Protease Inhibitor Cocktail (Roche), 0.25 U/µl Benzonase (Merck) and 0.5 U/µl ReadyLyse Lysozyme (Lucigen) in B-PER (ThermoFisher)]. Samples were incubated at room temperature for 5 min. 50 µl of 2x SDS sample loading buffer supplemented with 3 % v/v 2-ME were added. Samples were incubated at 95°C for 5 min, and the equivalent of 0.1875 OD units of cells was loaded onto a gel and analysed by SDS-PAGE. Western blotting was performed using a Trans-Blot turbo (BioRad) with the corresponding Midi 0.2 µm PVDF transfer pack. Blots were run at 25 V and 2.5 A for 7 min (mixed molecular weight programme). Membranes were blocked in PBS + 5 % milk for 30-40 min, washed in PBS and incubated with anti-HA-Peroxidase antibody (Roche 12013819001) (1/1000 in PBST + 5 % milk) at room temperature for 1-1.5h. Membranes were washed thoroughly with PBS and developed using the ECL Prime Western blotting detection kit (Amersham/Cytiva). Western blots were imaged on a Gel DocTM XR+ (Bio-Rad). Presented images were contrast-adjusted for display purposes.

## ACKNOWLEDGEMENTS

We thank D. Bellini, F. Gorrec and M. Yu (MRC LMB) for many helpful discussions on crystallisation and help with synchrotron data collection; the staff at Diamond Light Source beamlines I03 and I04 for excellent technical support; G. Cannone, C. Savva and all members of the LMB EM facility for training and support; G. Cannone and Y. Liu for help with EM data collection; T. Darling and J. Grimmett for help with computing infrastructure; K. Liu for WGS library preparation; all members of the Löwe group for discussions and technical help (all MRC LMB). T.N. was supported by a Boehringer Ingelheim Fonds (BIF) PhD fellowship and a Vice-Chancellor’s Award by the Cambridge Commonwealth, European & International Trust. F.B. was supported by an EMBO Advanced fellowship (ALTF 605-2019). This work was funded by the Medical Research Council (U105184326 to J.L.) and the Wellcome Trust (202754/Z/16/Z to J.L.).

## AUTHOR CONTRIBUTIONS

T.N. performed protein purifications, electron microscopy, co-pelleting, *in vivo* cysteine cross-linking and strain characterisation. S.H.M. performed SPR, FP and FDS-AUC experiments. T.N., D.K.-C. and J.L. performed crystallisation and crystallography. S.M. and M.J.S. performed HDX-MS experiments. C.W.H.Y. and S.M.V.F. performed NMR experiments and assignments. L.F.H.F. designed the genome mutagenesis strategy under supervision of J.W.C., which was adapted by F.B. for combinatorial mutagenesis. T.N. and F.B. performed genome engineering. T.N. and L.F.H.F. performed next generation sequencing. J.L. supervised the study. T.N. and J.L. wrote the manuscript with contributions from all authors.

## DATA AVAILABILITY

Atomic coordinates have been deposited in the PDB with accession codes 7Q6D (*Escherichia coli* FtsA^1-405^), 7Q6G (*Xenorhabdus poinarii* FtsA^1-396^), 7Q6F (*Vibrio maritimus* FtsA^1-396^, antiparallel double filament) and 7Q6I (*Vibrio maritimus* FtsA^1-396^ and FtsN^1-29^, bent tetramers in antiparallel double filament arrangement).

---

**Supplementary Figure S1.**
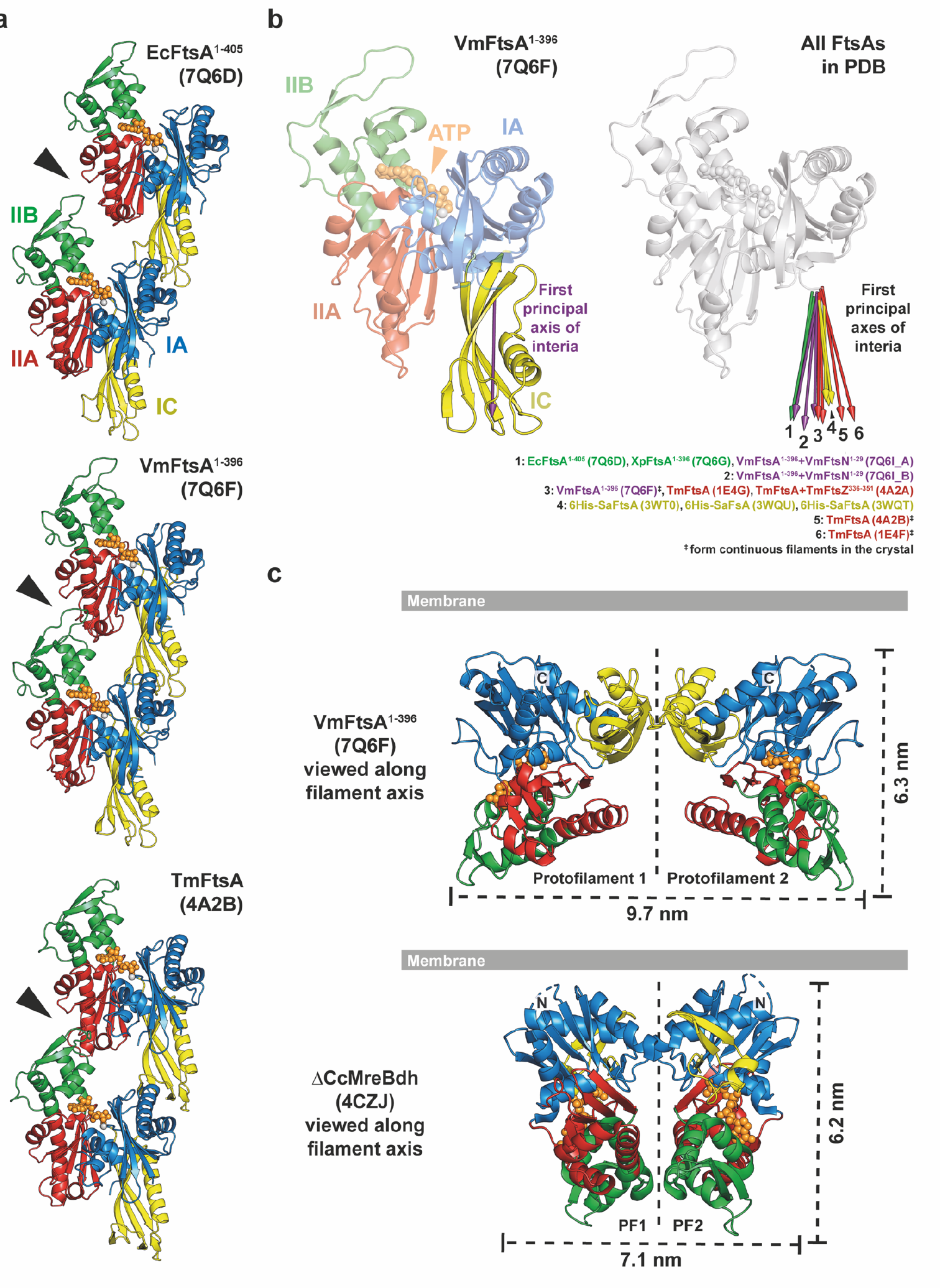
**a**, Comparison of longitudinal filament contacts in FtsA crystal structures from *E. coli* (PDB 7Q6D), *V. maritimus* (PDB 7Q6F) and *T. maritima* (PDB 4A2B). *E. coli* FtsA forms loose protofilaments with detached IIA and IIB domains (arrowhead). IIA and IIB domains are in close contact in the VmFtsA and TmFtsA structures, which form continuous filaments in the crystals. **b**, The IC domain of FtsA is flexible. Left: an arrow along the first principal axis of inertia of the IC domain (purple) can be used to indicate IC domain orientation. Principal axes of inertia were calculated using main chain atoms (N, CA, C) of IC domains. Right: FtsA structures in the PDB aligned on their IA, IIA and IIB domains, with arrows indicating the position of the IC domains, showing that the IC domain orientation is variable within the FtsA monomer. There is no correlation between IC domain position and species (different colours) or formation of continuous filaments in the crystals (^‡^). **c**, Comparison between the FtsA and MreB double filaments. Because the lateral interface is formed by the IC domain in the FtsA double filament, it is wider than the MreB double filament. The membrane-proximal side of both double filaments is flat.

**Supplementary Figure S2.**
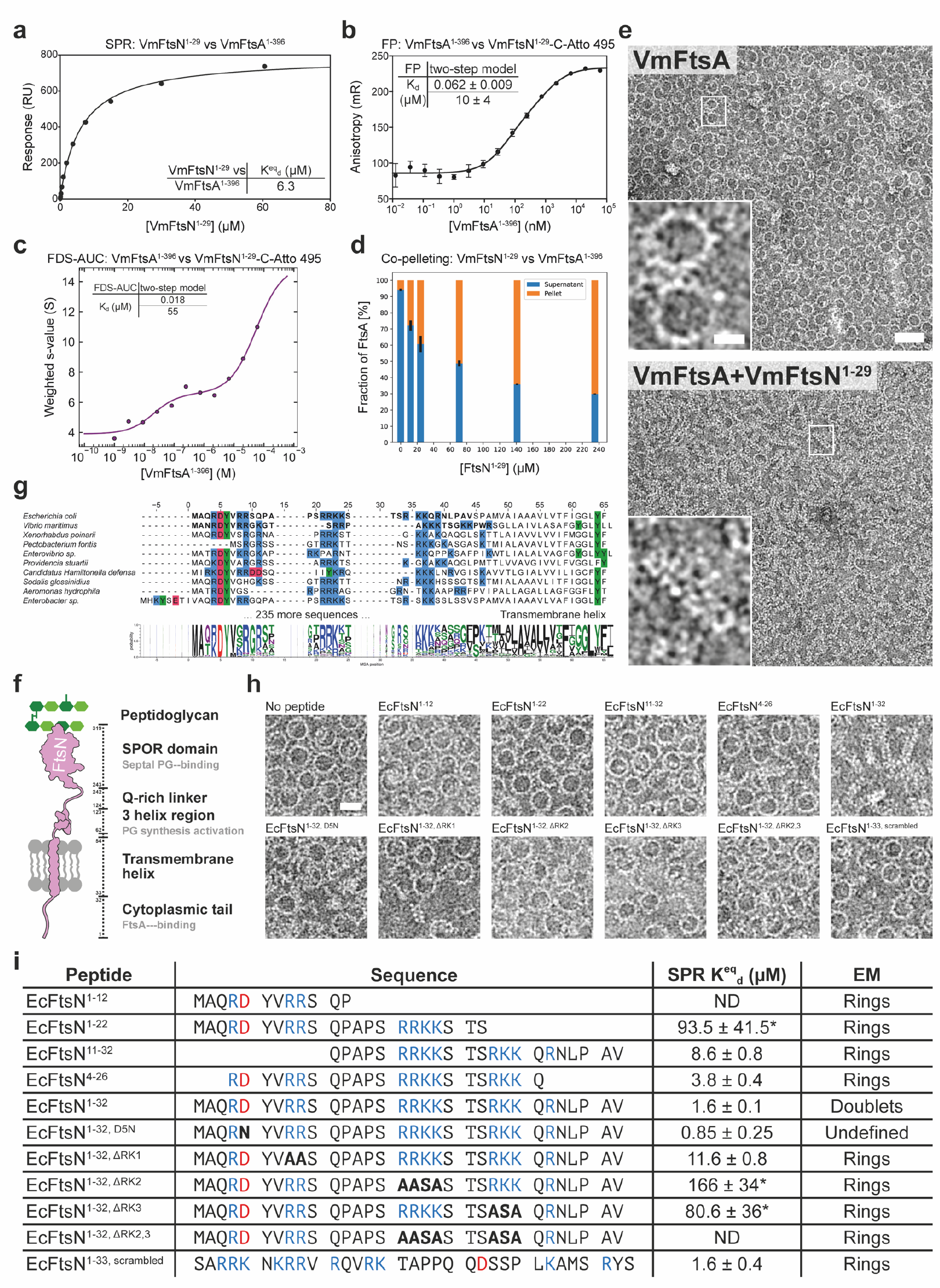
**a**, SPR equilibrium response titration of VmFtsN^1-29^ binding to immobilised VmFtsA^1-396^. Binding affinity is about three-fold lower than for the EcFtsA^1-405^-EcFtsN^1-32^ interaction (Figure 2a). **b**, VmFtsA^1-396^ titration into fluorescently labelled VmFtsN^1-29^-C-Atto 495. Data were fitted with a two-step model with transitions being indicative of FtsN binding and polymerisation (panel c). K_d_s are given as mean ± SEM. **c**, Change in the weight-averaged sedimentation coefficient of a VmFtsA^1-396^ titration into fluorescently labelled VmFtsN^1-29^-C-Atto 495 by FDS-AUC shows that VmFtsN^1-29^ is part of higher order FtsA polymers. Data were fitted to a two-step model, recapitulating the FP data in panel b. **d**, Co-pelleting assay of VmFtsN^1-29^ titrated into VmFtsA^1-396^. The fraction of FtsA in the pellet (orange) increases with increasing amounts of VmFtsN^1-29^, indicating that VmFtsN^1-29^ induces polymerisation of VmFtsA. Given are mean ± sd (black lines) of technical duplicates. **e**, Negative stain electron micrographs of VmFtsA with and without VmFtsN^1-29^ on supported lipid monolayers. Similar to *E. coli* FtsA, *V. maritimus* FtsA forms “mini-rings” in the absence of FtsN and double filaments at ten-fold molar excess of VmFtsN^1-29^. Scale bar, 50 nm, 20 nm (inset). **f**, Schematic overview of EcFtsN, illustrating domain boundaries. **g**, Multiple sequence alignment of 245 FtsN sequences comprising cytoplasmic and transmembrane domains. Ten exemplary sequences are shown including FtsN sequences from the organisms used in this study, namely *E. coli*, *V. maritimus* and *X. poinarii*. The sequences of EcFtsN^1-32^ and VmFtsN^1-29^ are highlighted in bold. **h**, Mapping of the FtsA-interacting region in EcFtsN^1-32^ using the lipid monolayer assay. EcFtsN^1-32^ truncations and mutants were in ten-fold molar excess of FtsA. In contrast to EcFtsN^1-32^, EcFtsN^1-32, D5N^ did not facilitate FtsA double filament formation. EcFtsN^4-26^ and EcFtsN^1-32, ΔRK1^ led to formation of fewer double filaments. A scrambled version of the FtsN peptide (Baranova *et al.*, 2020) did not induce double filaments. Scale bar, 20 nm. **i**, Summary of EcFtsN^1-32^ peptides. Mutations are highlighted in bold. Equilibrium dissociation constants (K^eq^) are given for SPR experiments as mean ± SEM (n ≥ 2 for each construct). For weak binders the maximum response was fixed during fitting, hence these are only approximate values as indicated by asterisks. “ND”, not determinable. The predominant higher order polymer observed in the monolayer assay is given in the “EM” column. Note that EcFtsN^4-26^ and EcFtsN^1-32, ΔRK1^ still lead to formation of a few FtsA double filaments. EcFtsN^1-33, scrambled^ matches a previously described version bearing an additional N-terminal cysteine and a C-terminal hexahistidine tag, which were removed for this study (Baranova *et al.*, 2020).

**Supplementary Figure S3.**
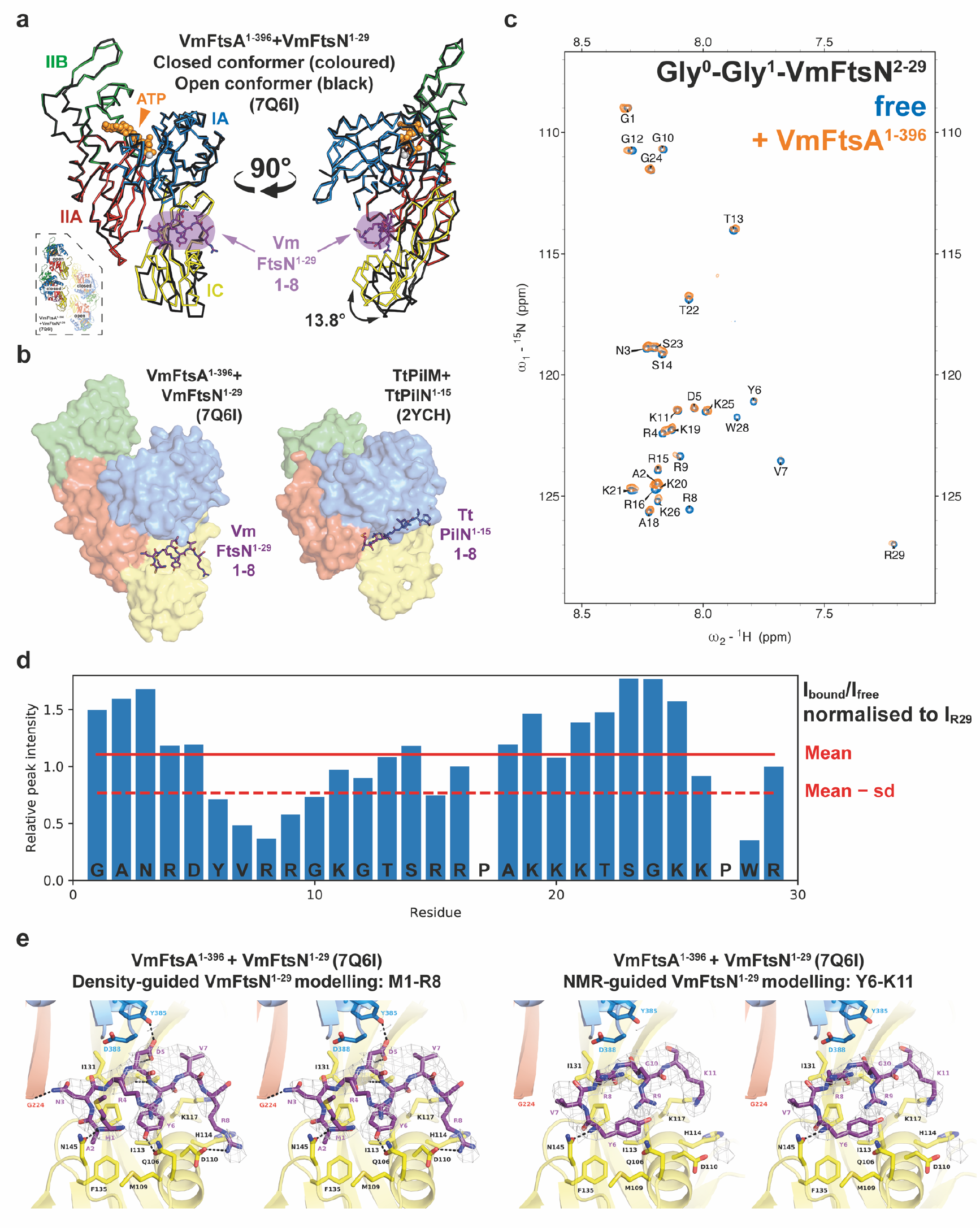
**a**, Comparison between the peptide-bound closed (coloured) and peptide-free open conformer (black) of FtsA in the VmFtsA^1-396^-VmFtsN^1-29^ co-crystal structure (PDB 7Q6I). The IC domain of the open conformer is rotated 13.8 ° downwards compared to the closed conformer, as determined by analysis with DynDom (Hayward and Lee, 2002). Consequently, the open conformation is likely incompatible with VmFtsN^1-29^ binding. The inset shows the position of open and closed conformers within the tetramer. **b**, Comparison between the *V. maritimus* FtsA-FtsN and *Thermus thermophilus* PilM-PilN (PDB 2YCH) interaction sites (Karuppiah and Derrick, 2011). While both binding sites are in the IA-IC interdomain cleft of FtsA and PilM, they occupy distinct subspaces. FtsN predominantly contacts the IC domain of FtsA, whereas PilN binds closer to the IA domain of PilM. **c**, ^1^H, ^15^N 2D-HSQC NMR spectrum of free Gly-Gly-VmFtsN^2-29^ (blue) and with equimolar amounts of FtsA added (orange). To follow VmFtsN numbering, the first glycine of Gly-Gly-VmFtsN^2-29^ is assigned as G0. **d**, Changes in relative peak intensity expressed as I_bound_/I_free_ with intensities normalised to I_R29_, which is assumed not to be involved in the VmFtsA-FtsN interaction. **e**, Stereo images of the FtsA-FtsN interaction site in the VmFtsA^1-396^-VmFtsN^1-29^ co-crystal structure (PDB 7Q6I). Left: our preferred interpretation of the electron density corresponding to VmFtsN^1-29^, with residues M1-R8 modelled (purple) and electron density map (grey) shown at 1.2 sigma. Side chains of FtsA residues in the interaction site are shown as sticks and polar contacts are marked with black, dashed lines. Right: electron density interpretation guided by the NMR data instead (panels c and d), with residues Y6-K11 of VmFtsN^1-29^ modelled (purple). Electron density map (grey) is shown at 1.2 sigma.

**Supplementary Figure S4.**
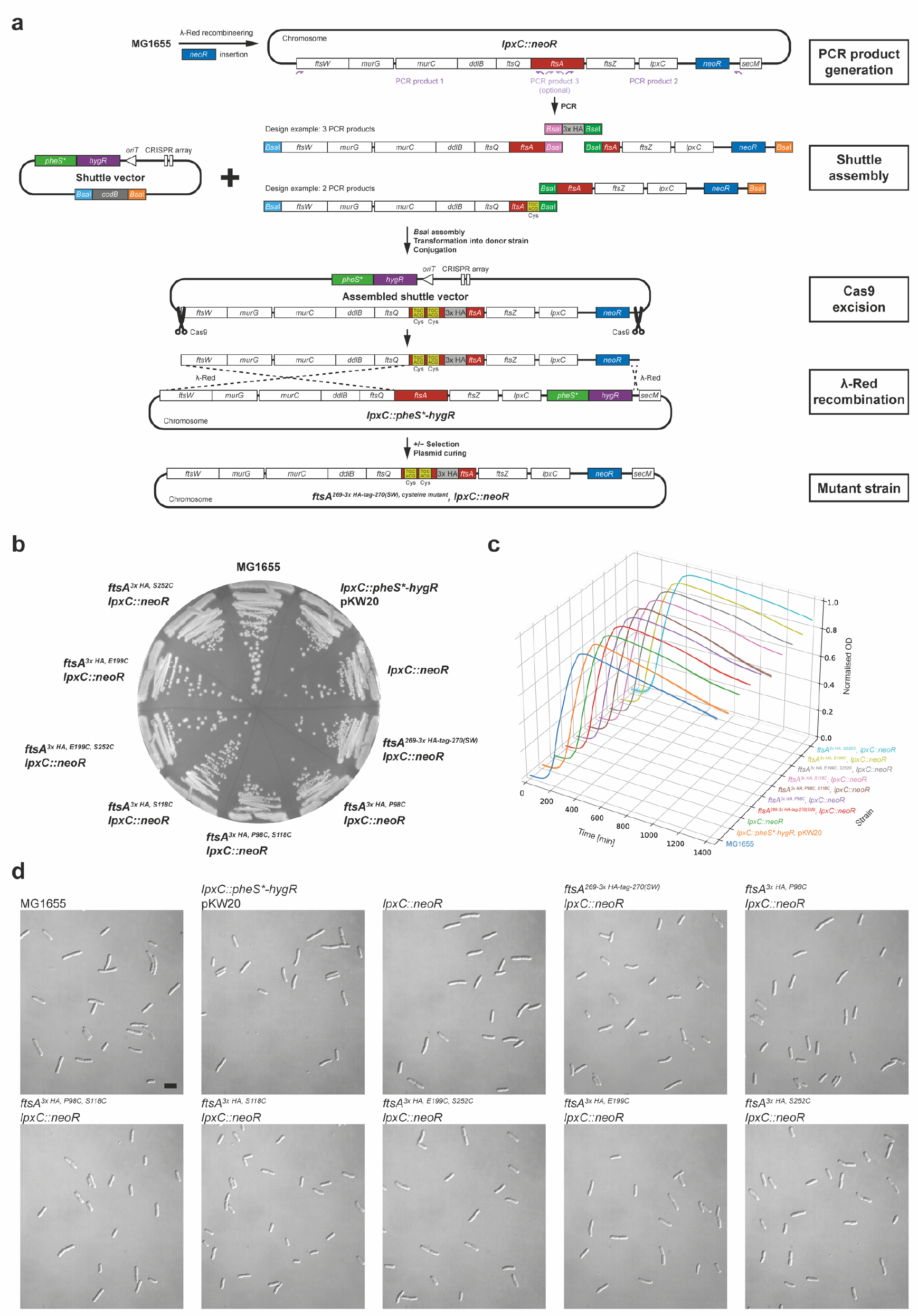
**a**, Workflow for REXER 2-based strain construction. PCR products containing the 3x HA-tag or cysteine point mutations were inserted into a shuttle vector by Golden Gate assembly (Engler *et al.*, 2008). Assembled shuttle vectors were transformed into the donor strain, conjugated, and excised *in vivo* using Cas9. Targeting constructs contain homology regions for *λ*-Red mediated recombination into the target locus. Recombinants were selected for *neoR* and *tetR* markers and against the *pheS\** marker. Strains were cured of the helper plasmid pKW20 by growth in absence of selection. **b**, Growth of strains containing single or double cysteine point mutations and a 3x HA-tag in the endogenous *ftsA* gene, and a kanamycin resistance cassette inserted after the *lpxC* gene. Parent strains and the original MG1655 strain are also shown. **c**, Growth curves of the same strains in liquid LB medium. Plotted are mean ± sd (technical octuplicates). **d**, DIC images of the same strains in exponential phase (OD_600_ = 0.2-0.3) demonstrating the absence of elongated cells.

## SUPPLEMENTS

**Supplementary Table T 1.**
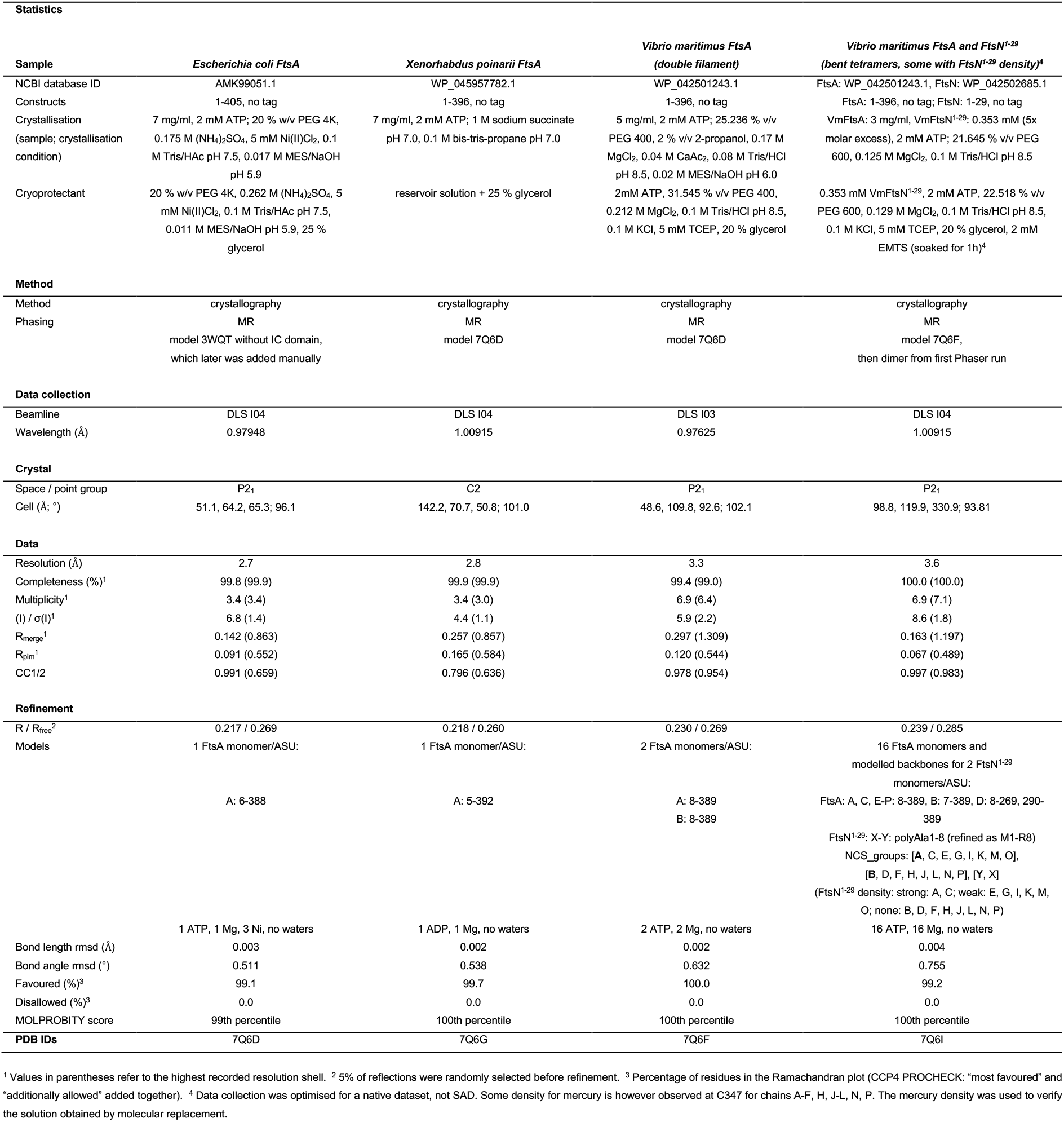
Crystallographic data.

**Supplementary Table T 2.**
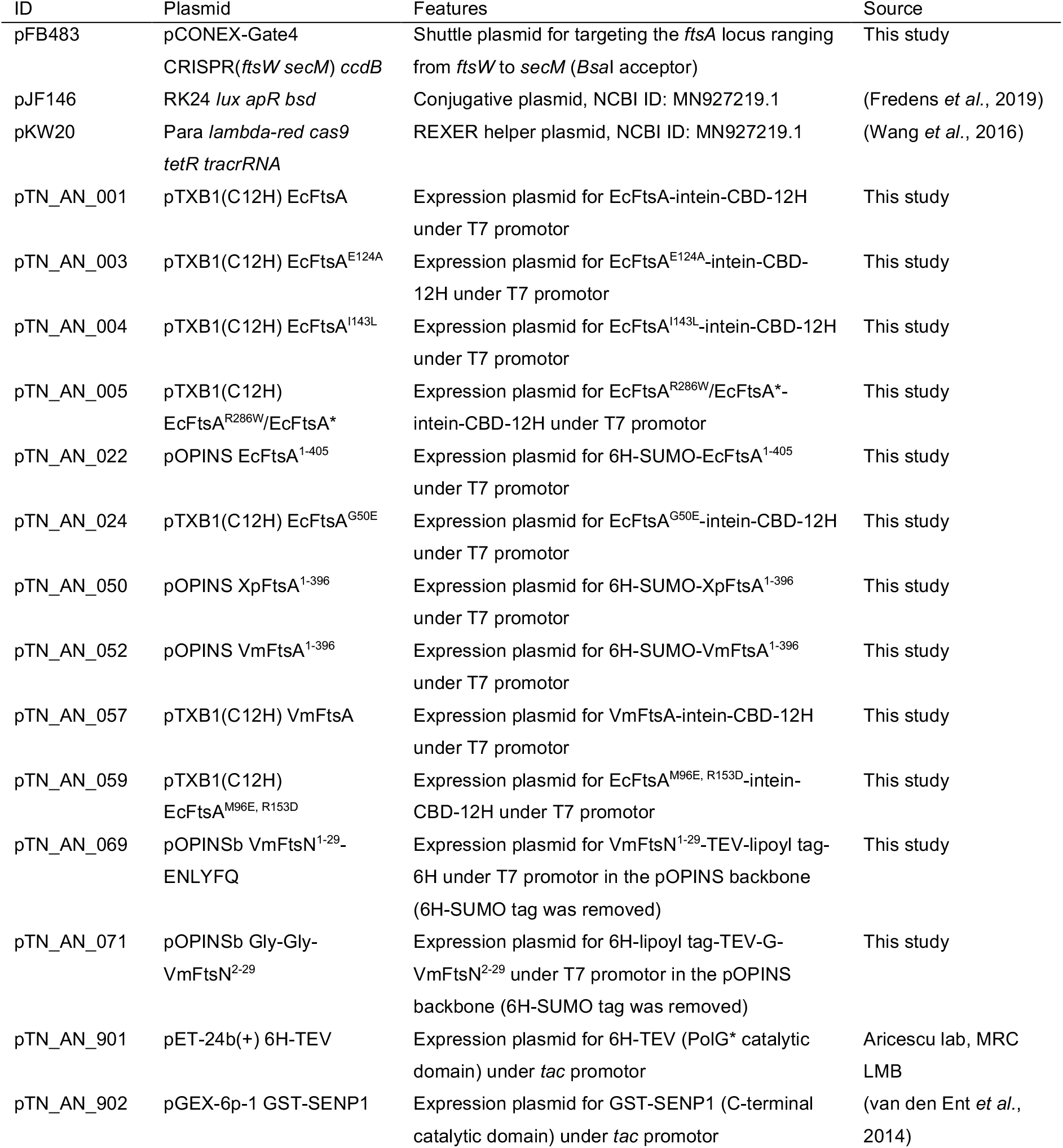
Plasmids.

**Supplementary Table T 3.**
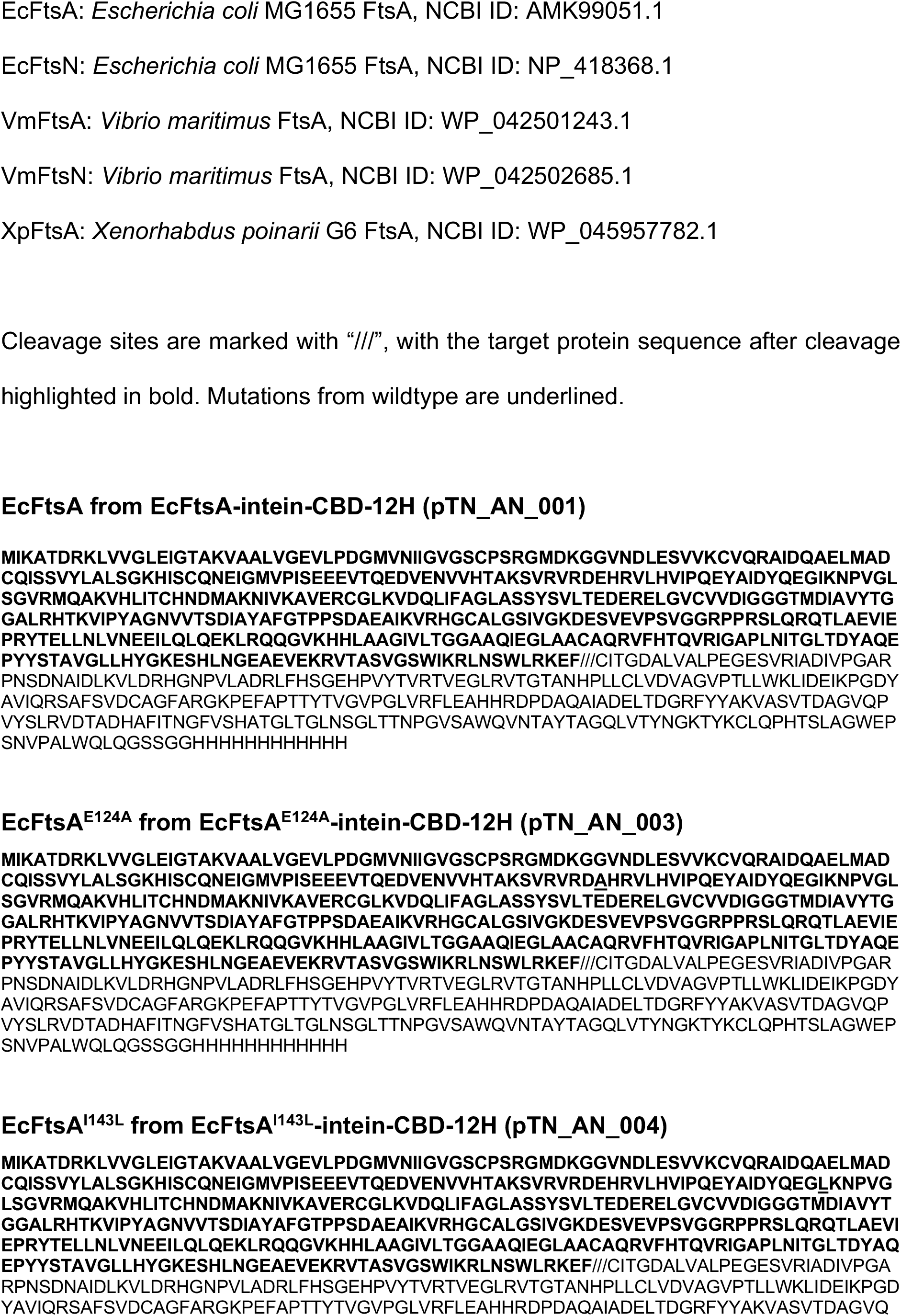
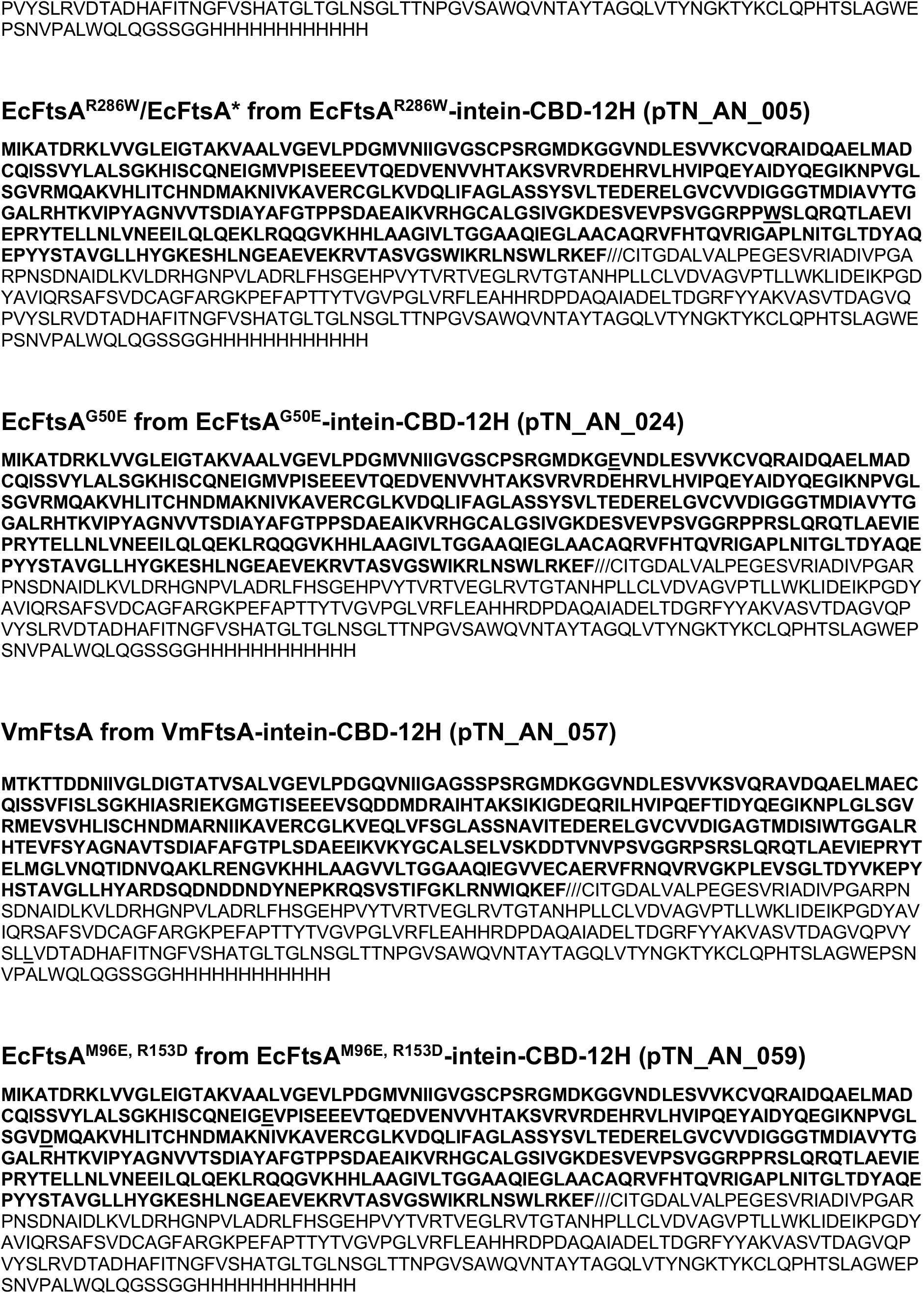
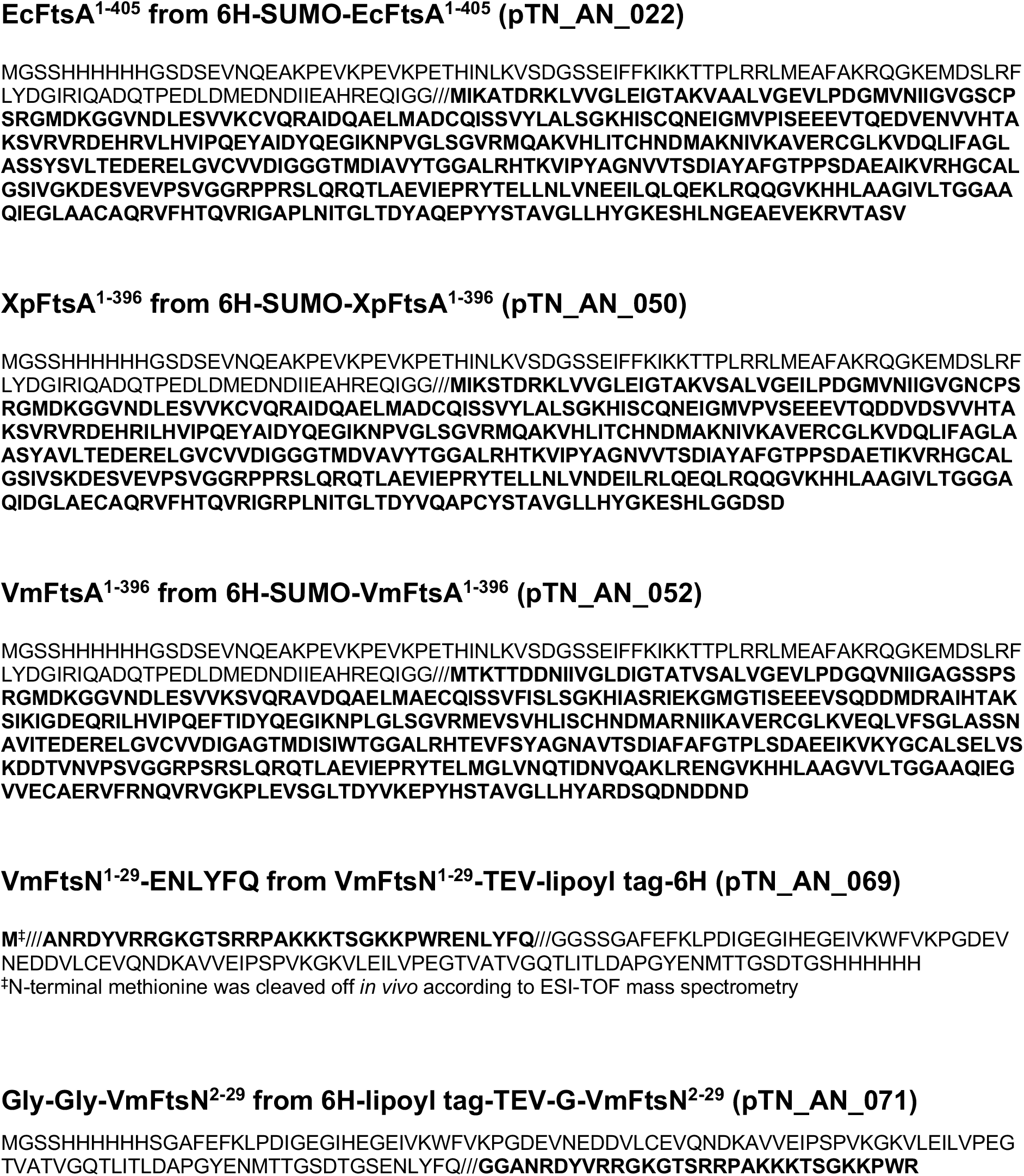
Protein sequences.

**Supplementary Table T 4.** FtsN peptide sequences.

| Peptide | Sequence | MW (Da) |
| --- | --- | --- |
| EcFtsN<sup>1-12</sup> | MAQ**RD** YV**RRS** QP | 1506.70 |
| EcFtsN<sup>1-22</sup> | MAQ**RD** YV**RRS** QPAPS **RRKKS** TS | 2605.96 |
| EcFtsN<sup>11-32</sup> | QPAPS **RRKKS** TS**RKK** Q**RNLP** AV | 2533.97 |
| EcFtsN<sup>4-26</sup> | **RD** YV**RRS** QPAPS **RRKKS** TS**RKK** Q | 2816.23 |
| EcFtsN<sup>1-32</sup> | MAQ**RD** YV**RRS** QPAPS **RRKKS** TS**RKK** Q**RNLP** AV | 3797.41 |
| EcFtsN<sup>1-32, D5N</sup> | MAQ**RN** YV**RRS** QPAPS **RRKKS** TS**RKK** Q**RNLP** AV | 3796.42 |
| EcFtsN<sup>1-32, ΔRK1</sup> | MAQ**RD** YV**AAS** QPAPS **RRKKS** TS**RKK** Q**RNLP** AV | 3627.19 |
| EcFtsN<sup>1-32, ΔRK2</sup> | MAQ**RD** YV**RRS** QPAPS **AASAS** TS**RKK** Q**RNLP** AV | 3529.00 |
| EcFtsN<sup>1-32, ΔRK3</sup> | MAQ**RD** YV**RRS** QPAPS **RRKKS** TS**ASA** Q**RNLP** AV | 3614.11 |
| EcFtsN<sup>1-32, ΔRK2,3</sup> | MAQ**RD** YV**RRS** QPAPS **AASAS** TS**ASA** Q**RNLP** AV | 3345.70 |
| EcFtsN<sup>1-33, scrambled</sup> | S**ARRK** N**KRRV** R**QVRK** TAPPQ Q**DSSP** L**KAMS** **RYS** | 3884.48 |
| EcFtsN<sup>1-32</sup>-C | MAQ**RD** YV**RRS** QPAPS **RRKKS** TS**RKK** Q**RNLP** AV**C** | 3900.55 |
| VmFtsN<sup>1-29</sup> | MAN**RD** YV**RRG** KGT**SR** RPA**KK** KTSG**K** KP**WR** | 3417.00 |
| VmFtsN<sup>1-29</sup>-C | MAN**RD** YV**RRG** KGT**SR** RPA**KK** KTSG**K** KP**WRC** | 3520.14 |

**Supplementary Table T 5.** *E. coli* strains.

| Strain | Genotype | Source | WGS* |
| --- | --- | --- | --- |
| *E. coli* | F<sup>-</sup> *ompT hsdSB (rB<sup>-</sup> mB<sup>-</sup>) gal dcm* (DE3); | Lucigen or Sigma | No |
| C41(DE3) | BL21(DE3) derivative |  |  |
| *E. coli* MAX | F<sup>-</sup> ϕ80*lacZ*ΔM15 Δ(*lacZYA-argF*) U169 *recA1* | ThermoFisher | No |
| Efficiency | *endA1 hsdR17* (rk<sup>-</sup>, mk<sup>+</sup>) *phoA supE44* λ<sup>-</sup> |  |  |
| DH5α | *thi-1 gyrA96 relA1* |  |  |
| MDS42 | MG1655, Δ*endA*, Δ*fhuABCD*, deletion of IS elements, prophages etc | Chin lab, MRC LMB (Posfai *et al.*, 2006) | No |
| MG1655 | F<sup>-</sup> λ<sup>-</sup> *rph-1 fnr<sup>+</sup>* | DSMZ (DSM 18039) | *rcIC*<sup>G169C</sup> |
| SFB123 | MG1655, *lpxC::pheS*<sup>T251A,A294G</sup>-*hygR*, pkW20<br>Para *lambda-red cas9 tetR tracrRNA* | This study | *rcIC*<sup>G169C</sup>,<br>*ppiD*<sup>D134G</sup> |
| SFB143 | MDS42, Δ*thiF*, pJF146 RK24 *lux apR bsd* | This study | No |
| SFB148 | MG1655, *lpxC::neoR*, pkW20 Para *lambda-red cas9 tetR tracrRNA* | This study | No |
| sTN001 | MG1655, *ftsA*<sup>269-3x HA-tag-270(SW)</sup>, *lpxC::neoR* | This study | *rcIC*<sup>G169C</sup>,<br>*ppiD*<sup>D134G</sup> |
| sTN003 | MG1655, *ftsA*<sup>P98C</sup>, 269-3x HA-tag-270(SW), *lpxC::neoR* | This study | *rcIC*<sup>G169C</sup>,<br>*ppiD*<sup>D134G</sup> |
| sTN005 | MG1655, *ftsA*<sup>P98C, S118C</sup>, 269-3x HA-tag-270(SW),<br>*lpxC::neoR* | This study | *rcIC*<sup>G169C</sup>,<br>*ppiD*<sup>D134G</sup> |
| sTN008 | MG1655, *ftsA*<sup>S118C</sup>, 269-3x HA-tag-270(SW), *lpxC::neoR* | This study | *rcIC*<sup>G169C</sup>,<br>*ppiD*<sup>D134G</sup> |
| sTN010 | MG1655, *ftsA*<sup>E199C, S252C</sup>, 269-3x HA-tag-270(SW),<br>*lpxC::neoR* | This study | *rcIC*<sup>G169C</sup>,<br>*ppiD*<sup>D134G</sup> |
| sTN011 | MG1655, *ftsA*<sup>E199C</sup>, 269-3x HA-tag-270(SW), *lpxC::neoR* | This study | *rcIC*<sup>G169C</sup>,<br>*ppiD*<sup>D134G</sup> |
| sTN014 | MG1655, *ftsA*<sup>S252C</sup>, 269-3x HA-tag-270(SW), *lpxC::neoR* | This study | *rcIC*<sup>G169C</sup>,<br>*ppiD*<sup>D134G</sup> |
| sTN015 | MG1655, *ftsA*<sup>Q155C</sup>, 269-3x HA-tag-270(SW), *lpxC::neoR* | This study | No |
| sTN017 | MG1655, *ftsA*<sup>D123C</sup>, 269-3x HA-tag-270(SW), *lpxC::neoR* | This study | No |

\* Intragenic, missense mutations only.

